# Targeting SAGA/ATAC complexes disrupts chromatin regulation and cholesterol metabolism in diffuse midline glioma

**DOI:** 10.64898/2026.01.22.701194

**Authors:** Rosemary U. Richard, Caitlin L. Bagnetto, Rebecca L. Murdaugh, Brittany R. Eberl, Alan L. Jiao, Adam F. Kebede, Ena I. Campos-Hensley, Barry M. Zee, Margaret M. Harrington, Mariella G. Filbin, Akdes Serin-Harmanci, Yang Shi, Jamie N. Anastas

## Abstract

Diffuse midline gliomas (DMG) are aggressive pediatric brain tumors characterized by chromatin and transcriptional dysregulation induced by H3K27M mutations, with a median survival of 11-15 months. We identified multiple components of the SAGA and ATAC chromatin regulatory complexes as DMG genetic dependencies and found that genetic or pharmacological inhibition of the SAGA/ATAC-associated chromatin reader SGF29 reduces DMG proliferation *in vitro* and prolongs survival in xenograft models. Small molecule inhibitors targeting SAGA/ATAC-associated histone acetylation, ubiquitination, and methylation similarly suppress DMG growth. Integrative chromatin and transcriptomic profiling reveals that disruption of SAGA/ATAC through *SGF29* knockout remodels the DMG chromatin landscape, producing distinct alterations at metabolic genes (loss of H3K9ac) and at embryonic/neurodevelopmental genes (redistribution of H3K4me3 and H3K27me3) and accompanying transcriptional changes. We further show that inhibition of the SAGA/ATAC-associated KAT2A/2B histone acetyltransferases represses cholesterol metabolism gene expression and that combined KAT2A/2B and cholesterol synthesis inhibition synergistically suppresses DMG growth *in vitro* and in a xenograft model. Together, these findings establish a mechanistic link between SAGA/ATAC-dependent chromatin regulation and the transcriptional and metabolic dysregulation underlying DMG malignancy.

**Significance:** We demonstrate that SAGA/ATAC-dependent chromatin regulation sustains malignant transcription controlling cholesterol metabolism, proliferation, and cell fate in DMG, suggesting that a SAGA/ATAC-regulated epigenome-metabolome axis may be targeted to treat DMG.

## Introduction

Somatic mutations resulting in a lysine 27 to methionine substitution in H3.1 or H3.3 (H3K27M) occur in ∼80% of cases of diffuse midline glioma (DMG) (1,2), which are near-universally fatal brain tumors diagnosed in children and adolescents (3). H3K27M mutations disrupt the balance of histone modifications by reducing repressive marks like H3K27me3/me2 while altering activating marks like histone acetylation and H3K4me3. These chromatin changes alter the expression of genes associated with mitosis, stemness, and progenitor states. (4–11). The enzymes and pathways driving oncogenic chromatin regulatory programs in DMG are not yet fully characterized.

The SAGA (Spt-Ada-Gcn5-acetyltransferase) chromatin regulatory complex is a conserved, multi-protein transcriptional co-activator that mediates cellular responses to environmental stressors and developmental cues (12–16). SAGA complexes are recruited to chromatin via scaffolding proteins like SGF29 (SAGA Complex Associated Factor 29), which binds H3K4-methylated histones found at enhancers and promoters (H3K4me3/2) (17), and TRRAP, which interacts with transcription factors linked to malignant transformation like MYC and CTNNB1 (18,19). Chromatin binding of SGF29 and TRRAP scaffolding proteins leads to the recruitment of various chromatin modifying enzymes, including lysine acetyltransferases like KAT2A and KAT2B and histone de-ubiquitinases (DUBs), which add acetyl groups and remove ubiquityl groups from chromatin, respectively (12,20–24). Alternatively, SGF29 can also recruit ATAC (ADA-two-A-containing) chromatin regulatory complexes, which similarly regulate histone acetylation through KAT2A and KAT2B and modulate H3K4me3 through WDR5-dependent recruitment of lysine methyltransferases (12,24–26). SAGA/ATAC complex genes are frequently overexpressed in tumors compared to normal tissue, and functional studies reveal that SAGA/ATAC-associated proteins promote disease progression in leukemia, breast cancer, prostate cancer, and glioma (13,27–36). Functional roles for SAGA/ATAC chromatin regulatory complexes in DMG are not yet established.

Epigenetic regulation and metabolism are tightly interconnected. Key cofactors and substrates required for histone acetylation, methylation, and ubiquitination are generated by metabolic pathways including glycolysis, the TCA cycle, fatty acid oxidation, and amino acid catabolism (37,38). Altered metabolism is a hallmark of glioma, with previous studies reporting aberrant mitochondrial activity, disrupted uptake and metabolism of lactate, glucose, and glutamine, and aberrant cholesterol synthesis in DMG (39–44). Cholesterol homeostasis and steroid metabolism are fundamental mediators of the growth and differentiation of oligodendrocyte progenitor cells (OPCs), thought to be the cell of origin for DMG (8,11,45), and represent promising therapeutic targets in glioma (33,37–40). However, direct inhibition of cholesterol biosynthesis may pose significant risks in pediatric patients, given the essential role of cholesterol in brain development and function. An alternative strategy for targeting DMG metabolic dysregulation might instead exploit DMG-specific chromatin and transcriptional regulation to decrease the expression of rate-limiting metabolic genes and inhibit tumor growth.

We identified multiple SAGA/ATAC complex genes as DMG genetic dependencies in a chromatin-focused CRISPR dropout screen. Complementary functional assays reveal that genetic or pharmacological targeting of SAGA/ATAC-associated chromatin modifying modules impairs DMG growth and that *SGF29* knockout slows DMG disease progression in orthotopic xenograft models. We conducted genome-wide chromatin and transcriptome profiling analyses, revealing that SAGA/ATAC inhibition alters DMG chromatin states at regions bound by H3K27M oncohistones and modulates the expression of genes linked to proliferation, differentiation, and cholesterol metabolism. Further studies suggest that SAGA-associated KAT2A/2B acetyltransferases promote DMG growth by inducing the transcription of cholesterol metabolism enzymes and reveal that co-targeting KAT2A/2B and cholesterol metabolism reduces DMG growth *in vitro* and *in vivo*.

Overall, these studies define roles for SAGA/ATAC complexes in sustaining aberrant gene regulatory programs in H3K27M mutant DMG, suggesting that SAGA/ATAC-dependent chromatin regulation constitutes a targetable mechanism underlying tumor maintenance and progression

## Materials and Methods

### Cell lines and culture conditions

Patient-derived H3K27M DMG cell lines (SU-DIPGVI, RRID:CVCL_IT40; SU-DIPGIV, RRID:CVCL_IT39; SU-DIPGXIII, RRID:CVCL_IT41; BT245, RRID:CVCL_IP13; and BT869, RRID:CVCL_E6HI cells) and H3 wildtype pHGG (SF9012, RRID:CVCL_E5GA; SF9402, RRID:CVCL_E5GB; SF9427, RRID:CVCL_E5GC) were kindly provided by Drs. Michele Monje, Keith Ligon, and Rintaro Hashizume. DMG cells were cultured in suspension as gliomaspheres in serum-free media DMG/HGG media consisting of a, 50:50 mix of DMEM/F-12 (cat#10-092-CV, Corning) and Neurobasal-A (cat#10888022, Gibco), 1x B27 minus Vitamin A (cat#12587010, Thermo), 1 mM Sodium Pyruvate (cat#25000CI, Corning), 2 mM GlutaMAX (cat#35050079, Gibco), 1x MEM non-essential amino acids (cat#11140050, Gibco), 0.01 M HEPES (cat#25-060-CI, Corning), and Antibiotic-Antimycotic (cat#30-004-CI, Corning). This DMG/HGG media also contained 0.002% Heparin (cat#07980, StemCell Technologies), hPDGF-AA (20 ng/mL; cat#10016100UG, Fisher Scientific), hPDGF-BB (20 ng/mL; cat#1001810, Fisher Scientific), hEGF (20 ng/mL; cat#100261MG, Fisher Scientific), and hFGF154 (20 ng/mL; cat#100146500UG, Fujifilm Irvine Scientific Inc). H3 wildtype DMG/HGG cells were maintained in adherent culture in DMG/HGG media supplemented with 5% FBS (cat#A5670701, Fisher Scientific). Cells were maintained by passaging every 4-6 days using TrypLE Express (cat#12604021, ThermoFisher). Adherent control cell lines (NHA-hTERT, RRID:CVCL_365 and BJ-Fibroblasts, RRID:CVCL_3653) and HEK293T (RRID:CVCL_0063) were purchased from ATCC and cultured in DMEM (cat#MT10013CV, Corning) containing 10% FBS and 1% Penicillin/Streptomycin (cat#SV3007901, Fisher Scientific). Adherent cells were passaged using 0.25% trypsin/EDTA (cat#MT25053Cl, Fisher Scientific). All cell lines were grown in a 5% CO2 incubator at 37C. Cell lines were validated by STR testing and checked regularly for mycoplasma contamination.

### CRISPR drop-out screen

SU-DIPGVI and SU-DIPGXIII cells expressing Cas9 were infected with a chromatin-focused sgRNA lentiviral library at a coverage of 1,500 cells per sgRNA in triplicate. Two days after infection, cells were selected for 3 days with 2 μg/mL puromycin (cat#P-600, Gold Biotechnology) and then allowed to grow for an additional three days. Reference samples were collected to quantify the initial sgRNA abundance. The remaining infected cells were outgrown for 5 weeks, and surviving cells were harvested. Genomic DNA was isolated from all samples and the sgRNA sequences were amplified by PCR before sequencing on a HiSeq2500 sequencer (Illumina). STARS was used to score depleted genes under each condition relative to the reference samples (https://portals.broadinstitute.org/gpp/public/software/stars).

### CRISPR sgRNAs and shRNAs

Oligonucleotide sequences targeting proteins of interest were designed using CRISPick (CRISPick), obtained from Millipore Sigma and phosphorylated with T4 polynucleotide kinase (cat#M0201L, NEB) before being cloned into pLenti SpBsmBI sgRNA hygro (cat#62205, AddGene, RRID:Addgene_62205) digested with Esp3I (cat#R0734L, NEB) using T4 DNA ligase (cat# M0202L, NEB). Plasmid products were cleaned using the NEB Monarch DNA gel extraction kit (cat#T1020S, NEB) and transformed into Stbl3 E. coli cells (cat#C737303, ThermoFisher, RRID: SCR_021177). A single colony was selected from each plate and plasmids were purified by mini-prep (cat#T1110L, NEB). DNA was quantified on a NanoDrop ND-1000 spectrophotometer and validated by Sanger sequencing. Sequence-validated plasmids were amplified in Stbl3 cells and purified using a Machery-Nagel Midiprep kit (cat#740420.50). As a control sgRNA for puromycin-containing plasmids, we used lentiCRISPR v2-sgControl (Addgene, RRID:Addgene_125836). KAT2A sgRNA1 and KAT2B sgRNA1 were obtained from AddGene (RRID:Addgene_138185, RRID:Addgene_138186). For clonal KO of *SGF29*, Lenti-iCas9-neo was used (AddGene, RRID:DGRC_8544) in combination with *SGF29*-targeting sgRNAs. The pLKO.1-puro plasmid (RRID:Addgene_8453) containing short hairpin RNAs (shRNAs) targeting *SGF29*, *KAT2A* and *KAT2B* were purchased from MilliporeSigma (KAT2Ash#1: TRCN0000038879, *KAT2*Ash#2: TRCN0000286981, *KAT2B*sh#1: TRCN0000018528, *KAT2B*sh#2: TRCN0000018529, *SGF29*sh#1: TRCN0000122343, *SGF29*sh#2: TRCN0000122343). A complete list of sgRNAs and shRNAs is provided in the Supplementary Table 1.

### Lentiviral transduction

Plasmids containing sgRNA and shRNA were co-transfected alongside packaging plasmids pCMV-dR8.2-dvpr (Addgene, RRID:Addgene_8455) and pCMV-VSVG (Addgene, RRID:Addgene_8454) into HEK293T (RRID:CVCL_0063) cells using PEI (cat#764965-1G, MilliporeSigma). Viral supernatants were collected at 48 and 72 hours post-transfection, pooled, and incubated with 50% PEG8000 (cat#BP233-1, Fisher Scientific) overnight at 4C. Supernatants were concentrated at 960G for 20 minutes and re-suspended in 600-900 uL of media per 10 cm plate. For lentiviral infection, 100-200 µl concentrated lentivirus was combined with 0.5×106-1×106 cells in 6-8xwells of a 12-well plate. Cells and virus were centrifuged at 960G for 90 minutes at room temperature and transferred to cell culture flasks to recover overnight in a 5% CO2, 37C incubator. 48-72 hours later, cells were selected with puromycin (cat#sc-108071A, SantaCruz Biotechnology), G418 disulfate salt (neomycin, cat#G8168, ThermoFisher), or hygromycin (cat#30240CR, Corning) as appropriate. For clonal growth assays, SU-DIPGXIIIP*+ZsG/luc cells were infected with Lenti-iCas9-neo and with Control sg, *SGF29*sgB, or *SGF29*sgC. Cells were removed from neomycin (cat#G8168, MilliporeSigma) selection at day 7 and allowed to recover for 4 days before disassociation and plating as single cells in 96-well plates by FACS. *SGF29*KO clones were validated by western blotting.

### Analysis of patient proteomics and RNA-seq datasets

Pediatric glioma proteomic and RNA-seq data were sourced from PedCBio portal (https://pedcbioportal.org/). Pediatric brain tumor samples were assigned into different categories based on the expression of proteins and mRNAs of interest, limiting these analyses to primary tumors or diagnostic samples. If multiple primary tumors or diagnostic samples from an individual patient were available, an average transcript or protein expression value was used in the analysis. High and low RNA expression groups were determined based on quartiles. High and low protein expression values were determined relative to the median protein expression value. Overall survival was analyzed using Kaplan-Meier curves generated in GraphPad Prism (RRID: SCR_002798) and survival differences between groups were determined using the Log-rank test.

### Cell proliferation and viability assays

Cell proliferation assays were performed by seeding 1×10^5^-5×10^5^ cells in 6-well plates and incubating with inhibitors for 24-120 hours. Treated cells were counterstained with 0.02% Trypan Blue and live/dead cells were counted using a hemocytometer. For cell viability assays, DMG and control cells were seeded at 4,000-10,000 cells per well in 96-well plates and treated with dose curves of various compounds. After 7 days, viability was quantified using the CellTiter-Glo Luminescent Cell Viability Assay (Promega, cat#G7573) according to the manufacturer’s protocol. Luminescence was measured on a PerkinElmer VICTOR X3 multilabel plate reader, and values were normalized to the DMSO-treated control condition.

### Western blot analysis

Whole cell lysates were prepared using RIPA lysis buffer (50 mM Tris-HCl pH 8.0, 0.1% SDS, 1% NP-40, 0.5% Na deoxycholate, and 150 mM NaCl) with protease (cat#5892970001, MilliporeSigma) and phosphatase inhibitors (cat#4906837001, MilliporeSigma) in 1x Laemmli buffer containing 2.5% v/v β-mercaptoethanol (cat#97064-878, VWR). Lysates were boiled at 95°C for 5 minutes, resolved on SDS-PAGE gels, and transferred to nitrocellulose (cat#10600001, Cytiva). Nitrocellulose membranes were then blocked in 4% non-fat dry milk in phosphate-buffered saline with Tween-20 (PBST) and incubated overnight with the following antibodies: total H3 (1:500,000; cat#ab1791, Abcam, RRID:AB_302613), H3K9ac (1:3,000, cat#9649, Cell Signaling Technology, RRID:AB_823528), H3K27ac (1:2,000, cat#39133, Active Motif, RRID:AB_2561016), H2AK119ub (1:2,000, cat#05-678, MilliporeSigma, RRID:AB_1080635), H3K27me3 (1:1,000, cat#07-449, MilliporeSigma, RRID:AB_310624), *SGF29 (1:2,000, cat#24061-1-AP, Proteintech, RRID:AB_2879420), GCN5L2 (1:2,000, KAT2A, cat#3305, Cell Signaling, RRID:AB_2128281), KAT2B (PCAF clone E.107.0, 1:1,000, cat#MA5-11186, ThermoFisher, RRID:AB_10978155), and USP22 (1:2,000, cat#sc-390585, Santa Cruz Biotechnology). The next day, membranes were washed with PBST, and incubated with 1:10,000-1:30,000 anti-rabbit (cat#B40962, ThermoFisher, RRID:AB_3096294) or anti-mouse HRP secondary antibodies (cat#B40961, ThermoFisher, RRID:AB_3096292) in 4% milk in PBST at room temperature. Membranes were imaged on a Bio-Rad ChemiDoc using ECL (cat#A38555, ThermoFisher).

### Immunofluorescence staining

To quantify H3K9ac levels, 250,000 BT869 cells were seeded onto laminin-coated coverslips and treated with vehicle (DMSO) or 25 μM SGF29-IN-1. Cells were fixed the following day with 4% paraformaldehyde in PBS for 20 min, washed extensively with PBST, and stored overnight at 4°C. The next day, coverslips were incubated in blocking buffer (5% FBS, 0.3% Triton X-100 in 1× PBS) for 45 min at room temperature. H3K9ac primary antibody (1:3,000; cat#9649, Cell Signaling, RRID:AB_823528) was diluted in primary antibody dilution buffer (0.3% Triton X-100, 1% BSA in 1× PBS) and incubated overnight at room temperature. The following day, coverslips were washed with PBST and incubated with Alexa Fluor 594 goat anti-rabbit IgG secondary antibody (1:1,000; cat#A11037, ThermoFisher, RRID:AB_2534095) for 30-45 min at room temperature. Coverslips were washed with PBST and stained with Hoechst 33342 (1:1,000; cat#51-17, Fisher Scientific) diluted in PBST for 10 min before mounting with Invitrogen ProLong Gold Antifade Mountant (cat#P36930, ThermoFisher). Images were acquired using a Keyence BZ-X microscope and analyzed using ImageJ (RRID:SCR_003070).

### Pharmacological inhibition of chromatin and cholesterol regulators

Small molecule inhibitors targeting various SAGA/ATAC-associated proteins or cholesterol synthesis enzymes were re-suspended in DMSO. The following compounds were used in this study: SGF29-IN-1 (cat# HY-158009, MedChemExpress), GSK4027 (cat#SML2018, MilliporeSigma), CPTH2 (cat#J65939LB0, ThermoFisher), garcinol (cat#14076, Active Motif), USP22i-S02 (cat#SML3875, MilliporeSigma), WDR5-IN-4 (WIN site inhibitor, cat#HY-111753, MedChemExpress), WDR5-IN-6 (WBM site inhibitor, cat#T77495, TargetMol), OICR9429 (cat#T6916, TargetMol), WM586 (WDR5/Myc inhibitor cat#HY-153728, MedChemExpress), YM-53601 (cat#HY-100313A, MedChemExpress), NB598 (cat#HY-16343, MedChemExpress), terbinafine (cat#T3677, TCI Chemicals), atorvastatin (cat#SML3030, MilliporeSigma), and pitavastatin (cat#AMBH2D6EF677, Ambeed). In selected studies DMG media was supplemented with water soluble cholesterol provided at 10 µM (cholesterol-methyl-β-cyclodextrin; cat#C4951, MilliporeSigma).

### Analysis of drug synergy

To calculate drug synergy, we set up dose curve matrices of pairwise drug combinations in various DMG cell lines and performed CellTiter-Glo viability assays after 7 days of treatment. We normalized the data to the average values for vehicle-treated cells. We determined drug synergy by BLISS analysis and generated heatmaps of synergy scores using the synergyfinder R package (version 3.12.0, RRID: SCR_019312) with the default settings. We also applied ZIP and Loewe scoring models in synergyfinder to confirm synergistic effects of different combinations of small molecules.

### Xenograft models of DMG tumorigenesis

NOD SCID gamma (NSG) mice were obtained from Jackson Labs (cat#005557, RRID:IMSR_JAX:005557). Orthotopic xenografts were established by transplant of 2.5×10^5^-5×10^5^ SU-DIPGXIIIP* or BT245 cells expressing Cas9, ZsGreen/luciferase and control or SGF29 sgRNAs into the pons of 6-8 week old mice at the following stereotactic coordinates zeroed on lambda: −1.0 mm X, −0.8 mm Y and −5.0 mm Z for pontine injections. Mice were assessed for weight loss, decline in body condition score, and neurological symptoms on a biweekly basis. *In vivo* monitoring of tumor size was performed with bioluminescence imaging (BLI). Briefly, mice were anesthetized with isoflurane and injected by IP with 150 mg/kg D-luciferin (cat#LUCK-5G, Gold Biotech). Eight minutes after injection, mice were imaged using the IVIS Lumina X5 Imaging system. After imaging, regions of interest were drawn around the head and spinal cords of mice and measurements of flux (p/s) were generated using Living Image software (v4.8.2, RRID: SCR_014247).

For studies of GSK4027 and pitavasatin activity *in vivo*, NSG mice (RRID:IMSR_JAX:005557) were injected with 2.5×105 DIPGXIIIP* cells directly into the pons. Two weeks post-surgery, mice received IP treatments four days a week for six weeks of vehicle (DMSO), pitavastatin (10 mg/kg), GSK4027 (10 mg/kg), or a combination of both drugs all prepared in sterile saline with 40% w/v PEG300 (cat#S6704, SelleckChem) and 0.1% v/v TWEEN80 (cat#P4780, MilliporeSigma). Mice were monitored weekly for tumor growth by bioluminescence imaging (BLI). All animal studies were performed according to the Baylor College of Medicine Institute Institutional Animal Care and Use Committee (IACUC)-approved protocols.

#### Brain tumor tissue collection, BrdU labelling, and IHC

After exposure to 3L/min CO2 for 3 minutes, mice were subjected to transcardial perfusion with saline followed by 4% paraformaldehyde (PFA). Brain tissue was dissected and fixed in 4% PFA overnight, followed by storage in 70% ethanol. For BrdU labeling *in vivo*, 24 hours after the final drug treatment, mice were pulsed with BrdU (100 mg/kg) for two hours. Paraffin embedding, hematoxylin and eosin (H&E), and BrdU staining (clone ZBU30; cat#03-3900, ThermoFisher, RRID:AB_2532917) was performed by the Smith Breast Center Pathology lab at Baylor College of Medicine using standard protocols on the Shandon Varistain (24–4) Automatic Stainer. Slides were imaged on a Keyence BZ-X800 microscope and the percentage of DAB+ BrdU stained cells within defined tumor regions was quantified using CellProfiler (RRID:SCR_007358).

### Histone mass spectrometry

Histone modifications were analyzed by mass spectrometry following acid extraction. DMG neurospheres were dissociated into single cells and incubated in swelling buffer (25 mM Tris-HCl pH 8.0, 1.5 mM MgCl₂, 10 mM KCl, 0.2% NP-40) supplemented with protease and phosphatase inhibitors and 10 mM sodium butyrate. Nuclei were collected by centrifugation at 464 RCF, washed in swelling buffer, and suspended in 0.2 N HCl containing butyrate and inhibitors. Histones were extracted overnight at 4°C with rotation. Acid extracts were cleared by centrifugation at 21,130 RCF for 30 min, and supernatants were transferred to fresh tubes. Trichloroacetic acid was then added to a final concentration of 30% (w/v) and proteins were precipitated on ice for 1 hour, followed by centrifugation at 21,130 RCF for 30 min. Pellets were washed with ice-cold 100% acetone, dried, and resuspended in 50 mM ammonium bicarbonate (pH 8.0). Sample quality was assessed by SDS-PAGE with Coomassie staining. - Histones were derivatized with propionic anhydride (1:3 propionic anhydride: isopropanol) before and after trypsin digestion (1:20 trypsin: histone, 50 mM ammonium bicarbonate, 4-6 h at 37°C). Histone peptides were desalted using in-house C18 STAGE tips and analyzed by liquid chromatography-mass spectrometry (LC-MS). Modified peptides were quantified by chromatographic peak integration as described previously (42).

### RNA-seq and CUT&RUN analysis

For SGF29KO RNA-seq experiments, 5×10^5^ control or *SGF29*KO BT245 and SU-DIPGXIII cells were collected in triplicate for processing. For RNA-seq following HATi treatment, 250,000 live cells were collected after 25 hours of treatment with DMSO, 10 µM GSK4027, or 10 µM CPTH2 in biological triplicates. RNA was extracted using TRIzol reagent (cat#15596018, Invitrogen) and submitted for paired-end RNA-seq through the Genomic and RNA Profiling Core (GARP) at Baylor College of Medicine. For bulk RNA-seq analysis, reads were aligned to hg38 genome build using STAR (RRID:SCR_004463) and read counts were determined on a gene level using featureCounts (RRID:SCR_012919). Differential gene expression analysis was performed using the Wald Test in DESeq2 (RRID:SCR_000154). Gene Ontology analysis was performed using the WEB-based GEne SeT AnaLysis Toolkit (RRID: SCR_011103 and GSEA analysis was performed using GSEA (v4.10, RRID: SCR_003199).

For CUT&RUN, a single-cell suspension of 5×10^5^ control or *SGF29*KO SU-DIPGXIII+Cas9 cells was immobilized on ConA beads, permeabilized with 0.01% digitonin, and incubated with the following primary antibodies: rabbit IgG (cat#C15410206, Diagenode, RRID:AB_2722554), H3K4me3 (cat#, C15410003 Diagenode, RRID:AB_2924768), H3K9ac (cat#9649, Cell Signaling, RRID:AB_823528), and H3K27me3 (cat#C15410195, Diagenode, RRID:AB_2753161). Yeast spike-in DNA was added at 0.5% to each reaction following chromatin digestion. Libraries were prepared from 3 ng of enriched DNA using the SMARTer ThruPLEX Plasma-seq Kit (cat#R400681, Takara Bio) with dual indexes (cat#DX34752, Takara Bio) according to the manufacturer’s instructions. Adapter-ligated libraries were amplified for 11 PCR cycles, purified using AMPure XP beads (cat#A63881, Beckman Coulter), and evaluated for fragment size distribution on an Agilent 2100 Bioanalyzer. Library molar concentrations were determined by quantitative PCR using the KAPA Library Quantification Kit (cat#KK4824, Roche) before pooling for sequencing on an Illumina NovaSeq 6000 to generate 150-bp paired-end reads. CUT&RUN sequencing data were aligned to hg38 using Bowtie2 after duplicate removal with Trim Galore (RRID:SCR_011847). MACS2 (RRID:SCR_013291) was used to call broad and narrow peaks, and consensus peak sets were analyzed with DiffBind (RRID:SCR_012918) to identify significantly upregulated and downregulated H3K4me3 and H3K9ac peaks in SGF29KO versus control cells. ChIP-seq data from previous studies generated using an H3K27ac antibody (cat#39133, Active Motif, RRID:AB_2561016), an H3K4me1 antibody (cat#ab8895, Abcam, RRID:AB_306847), and H3K27M CUT&RUN data generated using an H3.3 K27M mutant antibody (cat#ABE419, MilliporeSigma, RRID:AB_2728728) were also included in the integrative analysis (6,48). Motif analysis was performed using HOMER (RRID:SCR_010881) to identify enriched motifs within differential H3K9ac and H3K4me3 CUT&RUN peaks in SGF29KO SU-DIPGXIII cells. Input region size was set to 200 bp around peak centers, and repetitive sequences were masked. CUT&RUN peaks were also compared with previously generated transcription factor and chromatin regulatory protein ChIP-seq datasets using ChIP-Atlas (RRID:SCR_015511) to assess enrichment at known binding sites for gene regulatory factors.

### In vitro cholesterol assay

Cholesterol levels were determined using the Invitrogen Amplex Red Cholesterol Assay kit (cat#A12216, Invitrogen). BT969 cells plated at a density of 750,000 cells per well in 6-well plates were treated for 24 hours with either positive control inhibitors (FDFT1 or NB598) or with SAGA/ATAC-targeting compounds (SGF29-IN-1 and GSK4027). Lipids and cholesterol were extracted using the Folch method (2:1 chloroform: methanol) and analyzed in technical duplicates using the Amplex Red Cholesterol Assay kit (cat#A12216, ThermoFisher), processed according to manufacturer’s instructions, read using a fluorescence plate reader (FLUOstar Omega), and quantified according to a standard curve. Data presented are normalized to the DMSO-treated condition.

## Results

### SGF29, a SAGA/ATAC complex-associated chromatin reader, is required for optimal DMG growth

To identify chromatin regulators driving DMG tumorigenesis, we conducted CRISPR dropout screens in two independent H3.3K27M mutant DMG cell lines (SU-DIPGXIII and SU-DIPGVI cells) using an sgRNA library targeting 1,354 chromatin regulators with 6 sgRNAs per gene and hundreds of non-targeting guides as negative controls (6). STARS analysis (Screening-Through-Arrayed-Replicates System) of these data reveals a significant depletion in 149 sgRNAs (\**p*<0.01) in both SU-DIPGVI and SU-DIPGXIII cells (Supplementary Table 2). Further analysis of these CRISPR screens identifies multiple genes encoding members of the Spt-Ada-Gcn5 acetyltransferase (SAGA) complex and the related ADA-two-A-containing (ATAC) complex among the sgRNA dropouts (Fig. **1A** and **B**). These findings suggest that SAGA/ATAC chromatin regulatory complexes are required for optimal DMG growth *in vitro*.

**Figure 1.**
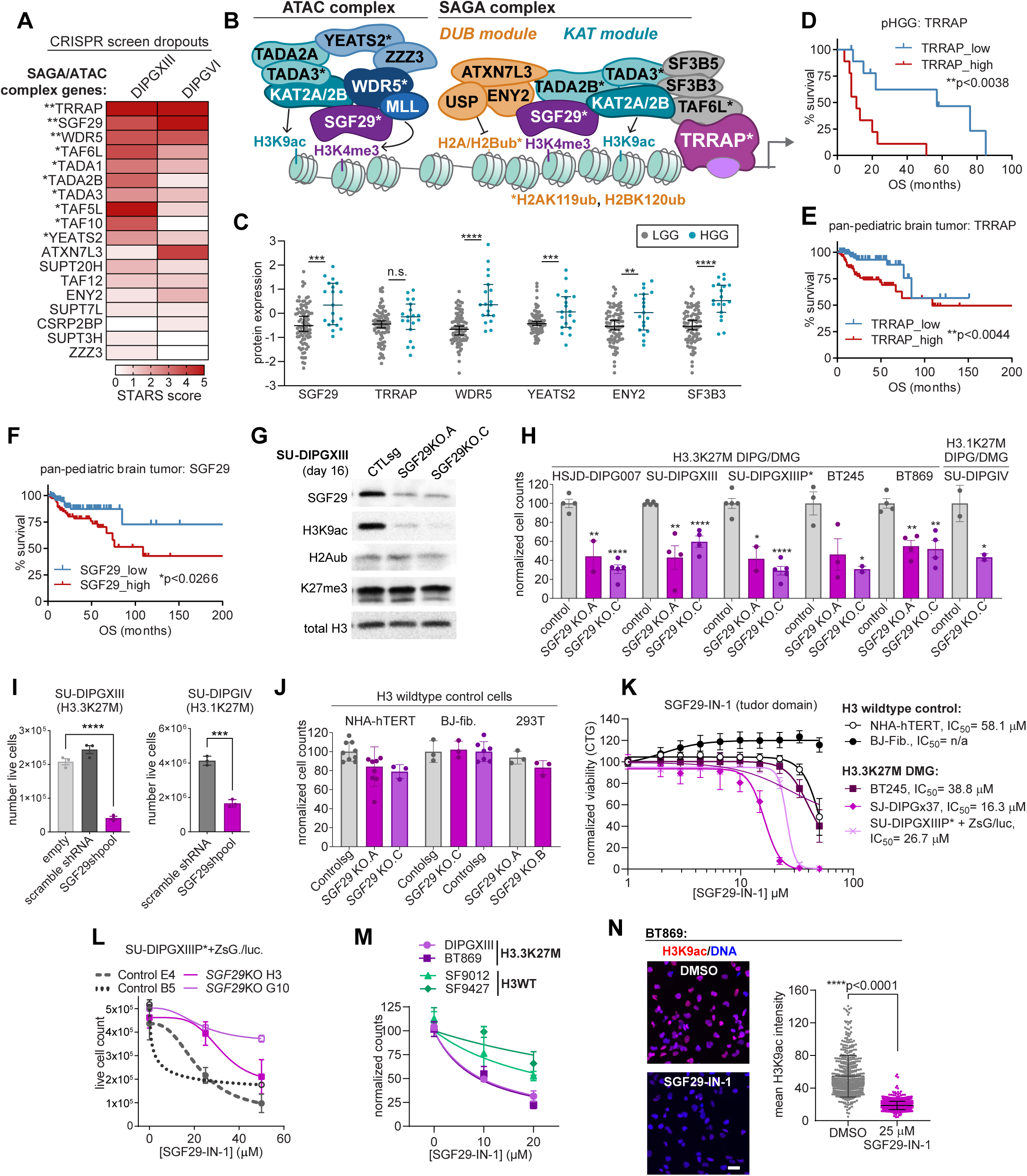
SGF29, a SAGA/ATAC complex-associated chromatin reader, is required for optimal DMG growth. **A**, Heatmap showing calculated STARS scores for SAGA/ATAC complex genes from a chromatin-focused CRISPR dropout screen, with red color correlating with stronger genetic dependencies and asterisks marking statistical significance. **B**, Schematic showing a subset of proteins found in the SAGA/ATAC chromatin regulatory complexes illustrating the H3K4me2/3 reader function of SGF29, KAT2A/2B-dependent histone acetylation, removal of H2AK119ub and H2BK120ub ubiquitination by the deubiquitnase (DUB) module, and ATAC-dependent regulation of H3K4me3. **C**, SAGA/ATAC-related protein expression in high grade (HGG) versus low grade glioma (LGG) primary pediatric brain tumor proteomics data (CPTAC). **D**-**F**, Kaplan-Meier survival curves comparing pediatric brain tumor patient outcomes grouped by low and high SGF29 and TRRAP protein expression in primary tumors. Panel **D** shows a survival analysis limited to pHGG, and panels **E** and **F** show pan-brain pediatric tumor analyses. **G**, Immunoblots of lysates from control and SU-DIPGXIII+Cas9 cells transduced with control or *SGF29* sgRNAs collected 16 days after starting selection showing a reduction in SGF29 protein abundance and SAGA/ATAC-targeted H3K9ac. **H**, Number of live Cas9(+) H3.3K27M and H3.1K27M mutant DMG cells 12-16 days after starting selection to express either control or *SGF29*-targeting sgRNAs. **I**, Number of live H3.3K27M mutant SU-DIPGXIII and H3.1K27M mutant SU-DIPGIV cells transduced with scramble (control) or pooled *SGF29* shRNAs. **J**, Number of non-transformed, H3 wildtype, astrocytes (NHA-hTERT), BJ fibroblasts (BJ fib.), or HEK293T cells following *SGF29*KO. **K**, CellTiter-Glo viability of DMG cells (purple curves) or H3 wildtype BJ fibroblasts and NHA-hTERT cells (black curves) following 7 days of treatment with increasing doses of SGF29-IN-1 to block SGF29 chromatin-binding via its tandem Tudor domains. **L**, Average number of control and clonal *SGF29*KO SU-DIPGXIIIP*+ZsG/luc cells confirming on-target inhibition of SGF29 with SGF29-IN-1. **M,** Dose curves showing that H3 wildtype pHGG are less sensitive to SGF29-IN-1 than H3K27M mutant DMG. **N**, Representative immunofluorescence images and quantification of mean H3K9ac intensity from BT869 cells treated with vehicle or 25 µM SGF29-IN-1 for 24 hours.

Specific SAGA/ATAC complex members identified as DMG genetic dependencies include the H3K4me3/2 chromatin reader, *SGF29* (SU-DIPGVI: *FDR*<0.001; SU-DIPGXIII: *FDR*<0.0062), the SAGA-associated scaffolding protein, *TRRAP* (SU-DIPGVI: *FDR*<0.00037; SU-DIPGXIII: *FDR*<6.48×10^-5^), and additional SAGA/ATAC core components (*TAF6L*, *TADA1*, *TADA2B*, *TADA3*) (Fig. **1A** and **B**). ATAC-associated chromatin regulators were also identified as DMG genetic dependencies including *WDR5*, which regulates H3K4me3 through MLL methyltransferase recruitment to ATAC and moonlights in other chromatin regulatory complexes like the COMPASS complex (49), and *YEATS2*, a core component of the ATAC-associated HAT module (**Fig. 1B**) (12,50). While a previous study identified KAT2A as a genetic dependency in H3.1K27M mutant DMG (36), SAGA/ATAC-associated HATs, *KAT2A* and *KAT2B,* were not identified as DMG genetic dependencies, potentially due to genetic compensation or functional redundancy as has been previously reported in some contexts (24,51).

We then analyzed patient RNA-seq and proteomics datasets from PedcBioPortal to assess correlations between SAGA/ATAC mRNA and protein expression and pediatric brain tumor patient outcomes. mRNAs encoding SAGA/ATAC components are expressed at elevated levels in pediatric high grade glioma (HGG) compared to low grade glioma (LGG) patient samples (Supplementary Fig. S1A) and a subset of SAGA/ATAC complex members were similarly overexpressed at a protein level in HGG versus LGG, including SGF29, WDR5, YEATS2, ENY2, and SF3B3 (**Fig. 1C**). Kaplan-Meier survival analyses comparing patients grouped by high (quartile 4), medium (quartiles 2-3), and low (quartile 1) *SGF29* and *TRRAP* mRNA expression in primary tumors/diagnostic samples suggest that increased expression of these core SAGA components is associated with poor patient outcomes in a pan-pediatric brain tumor cohort (Supplementary Fig. S1B and S1C). Increased TRRAP protein expression similarly correlated with decreased survival in both a pan-pediatric brain tumor cohort (\*\**p*<0.0044) and in an HGG patient cohort (\*\**p*<0.0038) (Fig. **1D** and **E**). Elevated SGF29 protein abundance similarly correlated with poor clinical outcomes in the pan-pediatric brain tumor cohort (\**p*<0.026) (**Fig. 1F**) but did not reach significance in HGG (*p*<0.0715, Supplementary Fig. **1D**). Further analyses indicate that increased expression of additional SAGA-associated proteins (TAF9, ENY2, SF3B3) and ATAC components (WDR5, YEATS2) similarly predicted worse patient outcomes in the pan-pediatric glioma proteomic dataset (Supplementary Fig. S1E-I). These results confirm that SAGA/ATAC-associated transcripts and proteins are detected in pediatric brain tumor patient samples and suggest that increased SAGA/ATAC protein and RNA expression correlates with reduced survival.

To assess whether SAGA/ATAC might represent a specific dependency in H3K27M mutant brain tumors, we queried the Broad pan-cancer dependency (DepMap) database and calculated average CHRONOS dependency scores across cancer subtypes arising in different tissues. We included a strong universal genetic dependency (*KIF11*) and two common tumor suppressors (*TP53*, *PTEN*) to capture the range of responses to different CRISPR knockouts (top and bottom rows of heatmap, Supplementary Fig. S1J). This analysis suggests that SAGA-associated *TRRAP* and *SF3B5* and *SF3B3* splicing regulators were strong dependencies across all cancer subtypes on par with *KIF11* (Supplementary Fig. S1J). In contrast, most of the SAGA/ATAC genes were categorized neither as strong dependencies nor as potential tumor suppressors in these adult cancers (Supplementary Fig. S1J). These analyses highlight SAGA/ATAC complexes as potentially unique drivers of H3K27M DMG malignancy, complementing previous studies linking SAGA/ATAC-associated proteins to disease progression in a subset of cancers like leukemia, breast cancer, prostate cancer, and adult brain cancer (13,27–36).

Based on our CRISPR screen results and our analyses indicating that increased SAGA/ATAC expression correlates with reduced survival in pediatric glioma patient datasets, we then assessed a potential role for SGF29 in promoting DMG growth. We selected SGF29 for these initial studies since it is a member of both the SAGA and ATAC complexes where it functions as a chromatin-binding scaffold to recruit SAGA/ATAC-associated chromatin modifying modules to gene targets (**Fig. 1B**) (17), and does not represent a universal, pan-cancer dependency (Supplementary Fig. S1J), suggesting some selectivity for DMG. We transduced stable Cas9-expressing H3.3K27M mutant DMG cell lines (HSJD-DIPG007, SU-DIPGXIII, SU-DIPGXIIIP*, BT245, and BT869) and an H3.1K27M mutant cell line (SU-DIPGIV) using lentiviral transduction to express either a non-targeting control sgRNA or *SGF29*-targeting sgRNAs to knockout (KO) *SGF29*. Immunoblotting of whole cell lysates extracted from polyclonal SU-DIPGXIII+Cas9 cells reveals a loss of SGF29 protein due to transduction with two independent sgRNAs (*SGF29*KO.A and *SGF29*KO.C) and a corresponding reduction in a known SAGA/ATAC-regulated histone acetyl mark (H3K9ac; **Fig. 1G**) (24), indicating effective genetic targeting of SAGA/ATAC-mediated histone acetylation. *SGF29KO* reduced the growth and survival of multiple H3.3K27M mutant cell lines and an H3.1K27M mutant cell line (SU-DIPGIV) in neurosphere culture (**Fig. 1H**) and reduced SU-DIPGXIIIP* growth as single-cell clones (Supplementary Fig. S1K) compared to control sgRNA-transduced cells. shRNA knockdown of *SGF29* similarly reduced both H3.3K27M (SU-DIPGXIII) and H3.1K27M (SU-DIPGIV) cell growth compared to scramble shRNA control (**Fig. 1I**). In contrast, *SGF29*KO did not significantly reduce the growth of H3 wildtype, non-transformed cell lines such as HEK293T, BJ fibroblasts, and immortalized astrocytes (NHA-hTERT) (**Fig. 1J**). To further confirm a role for SAGA/ATAC complexes in promoting DMG growth, we also generated *TRRAP*KO SU-DIPGXIII and BT869 cells and similarly observe reduced growth due to loss of this SAGA complex scaffold protein (Supplementary Fig. S1L and S1M).

To complement these SAGA/ATAC gene knockout studies, we determined the effect of treating H3K27M mutant DMG and control cells with SGF29-IN-1, a chemical probe recently developed to inhibit SGF29 binding to H3K4me2/3-modified chromatin by blocking its chromatin-reading Tudor domains (32). Dose curve analyses using CellTiter-Glo viability assays and live cell counts as endpoints reveal that multiple DMG cell lines were sensitive to pharmacological inhibition of SAGA/ATAC by SGF29-IN-1 treatment (Fig. **1K**-**N**), whereas non-transformed BJ fibroblasts and NHA-hTERT cells were not responsive up to a 50 µM dose (black curves, **Fig. 1K**). Additional studies reveal that *SGF29*KO clones exhibit reduced sensitivity to SGF29-IN-1 treatment, confirming on-target inhibition of SGF29 at these doses (**Fig. 1L**), and reveal that two H3 wildtype pediatric high grade glioma cell lines (pHGG; SF9012 and SF9427) were less sensitive than H3.3K27M cell lines to SGF29-IN-1 (**Fig. 1M**). Further analyses confirm that SGF29-IN-1 reduces H3K9ac staining intensity in BT869 cells, consistent with effective targeting of SAGA/ATAC-dependent histone acetylation (**Fig. 1N**). Together, these results demonstrate that either genetic or pharmacological inhibition of SGF29 selectively reduces the proliferation and survival of multiple H3K27M mutant DMG cell lines in culture compared to H3 wildtype HGG and control cells.

### SGF29 promotes DMG tumor growth and progression

We next asked whether SGF29 is required for DMG tumor growth in xenograft models. We transplanted control and clonal *SGF29*KO SU-DIPGXIIIP*+Cas9 cells co-expressing ZsGreen and luciferase (ZsG/luc) into the pons of immunocompromised NSG mice and monitored tumor growth by bioluminescent imaging (BLI) (Fig. **2A-C**). While the tumor BLI signal was not significantly different at 3 days post-transplant, confirming that *SGF29*KO did not impair initial engraftment (Fig. **2B**), further analyses reveal a significant decrease in BLI signal in the mice bearing *SGF29*KO xenograft tumors compared to control sgRNA tumors, suggesting reduced tumor size (Fig. **2A** and **C**). We confirmed the reduction in *SGF29*KO SU-DIPGXIIIP* tumor size compared to controls through histological analysis of post-mortem brains, revealing that the control xenograft tumors exhibited extensive growth and leptomeningeal invasion compared to the much smaller *SGF29*KO tumors (Fig. **2D**). Monitoring body weight as an overall indicator of health suggests that mice in the control sgRNA group exhibited rapid weight loss, while body weight and overall health were maintained over a prolonged period in the *SGF29*KO cohort, consistent with reduced disease burden (Fig. **2E**). A Kaplan-Meier analysis confirms a significant increase in the survival of the mice bearing *SGF29*KO versus control tumors (\*\*\**p*<0.0001, Log-Rank test; Fig. **2F**). We verified an *in vivo* dependency on *SGF29* in a second orthotopic xenograft model (BT245+Cas9+ZsG/luc), similarly demonstrating increased survival in the *SGF29*KO group (\**p*<0.0095, Log-rank test; Fig. **2G**). These findings implicate *SGF29* as a regulator of DMG tumorigenesis and suggest that SAGA/ATAC-dependent chromatin regulation might be targeted to reduce DMG disease progression *in vivo*.

**Figure 2.**
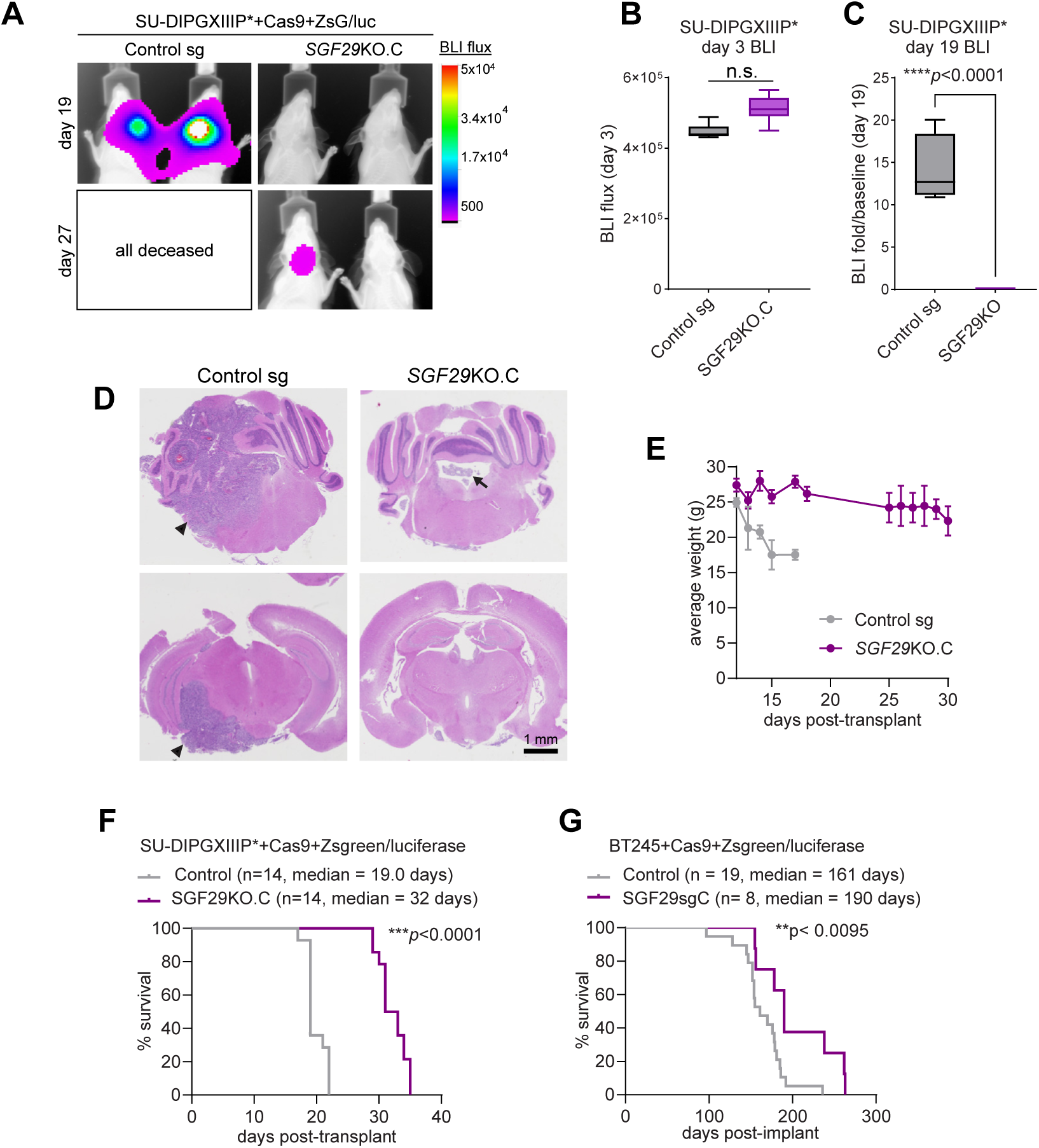
SGF29 promotes DMG tumor growth and progression**. A**-**C**, Representative bioluminescent (BLI) images (**A**) and quantification (**B** and **C**) of tumor luciferase signal in mice bearing orthotopic pontine xenografts of control and *SGF29*KO SU-DIPGXIIIP*+Cas9+ZsG/luc cells showing no significant difference in tumor luciferase signal at day 3 (**B**) but reduced *SGF29*KO DMG tumor luciferase normalized to baseline at day 19 post-transplant (**C**). **D**, Representative images of H&E-stained coronal brain sections from the SU-DIPGXIIIP* xenografts showing large tumor masses in the hindbrain and midbrain of a mouse transplanted with control cells (black arrowheads) and much smaller tumor mass in the 4^th^ ventricle of the *SGF29*KO tumor-bearing mouse (black arrow). **E**, Average weight (grams) of mice bearing SU-DIPGXIIIP* xenograft tumors indicating a prolonged maintenance of normal body weight in the *SGF29*KO cohort. **F** and **G**, Kaplan-Meier survival analysis of *SGF29*KO versus control xenograft cohorts suggesting that *SGF29*KO slows DMG disease progression in both the SU-DIPGXIIIP*+Cas9+ZsG/luc xenograft model (\*\*\**p*<0.0001, Log-rank test; **F**), and in the BT245+Cas9+ZsG/luc xenograft model (\**p*<0.0095, Log-rank test; **G**).

### Genetic and pharmacological inhibition of SAGA/ATAC-associated chromatin modifying modules inhibits DMG growth

A key function of SAGA/ATAC chromatin regulatory complexes is to activate RNA Polymerase 2 (Pol2)-dependent transcription of target genes via the recruitment of enzymes and accessory proteins that regulate histone post-translational modifications (PTMs) like acetylation, methylation, and ubiquitination (12,51). We next asked whether genetic or pharmacological targeting of SAGA/ATAC-associated histone modifying activities can regulate DMG proliferation *in vitro*. To genetically inhibit SAGA/ATAC-dependent histone acetyltransferase (HAT) activity, we transduced various Cas9-expressing DMG cell lines with *KAT2A* and *KAT2B* sgRNAs. While KO of either *KAT2A* or *KAT2B,* as validated by western blot, reduced the growth of SU-DIPGVI cells (Supplementary Fig. S2A and S2B), loss of these HATs had no significant effect on the growth of SU-DIPGXIIIP*, BT869, or SU-DIPGXIII cells (Supplementary Fig. S2C-S2E), potentially due to partially redundant roles for these enzymes in regulating histone acetylation. Indeed, co-infection with multiple shRNAs to simultaneously knockdown both *KAT2A* and *KAT2B* significantly reduced SU-DIPGXIII cell growth (Fig. **3A**). KO of *TADA2B*, which is required for KAT2A/2B recruitment to SAGA complexes (Fig. **1B**) (12,24), similarly reduced the growth of SU-DIPGXIII cells, offering additional evidence that the SAGA HAT module promotes DMG proliferation (Fig. **3B**).

**Figure 3.**
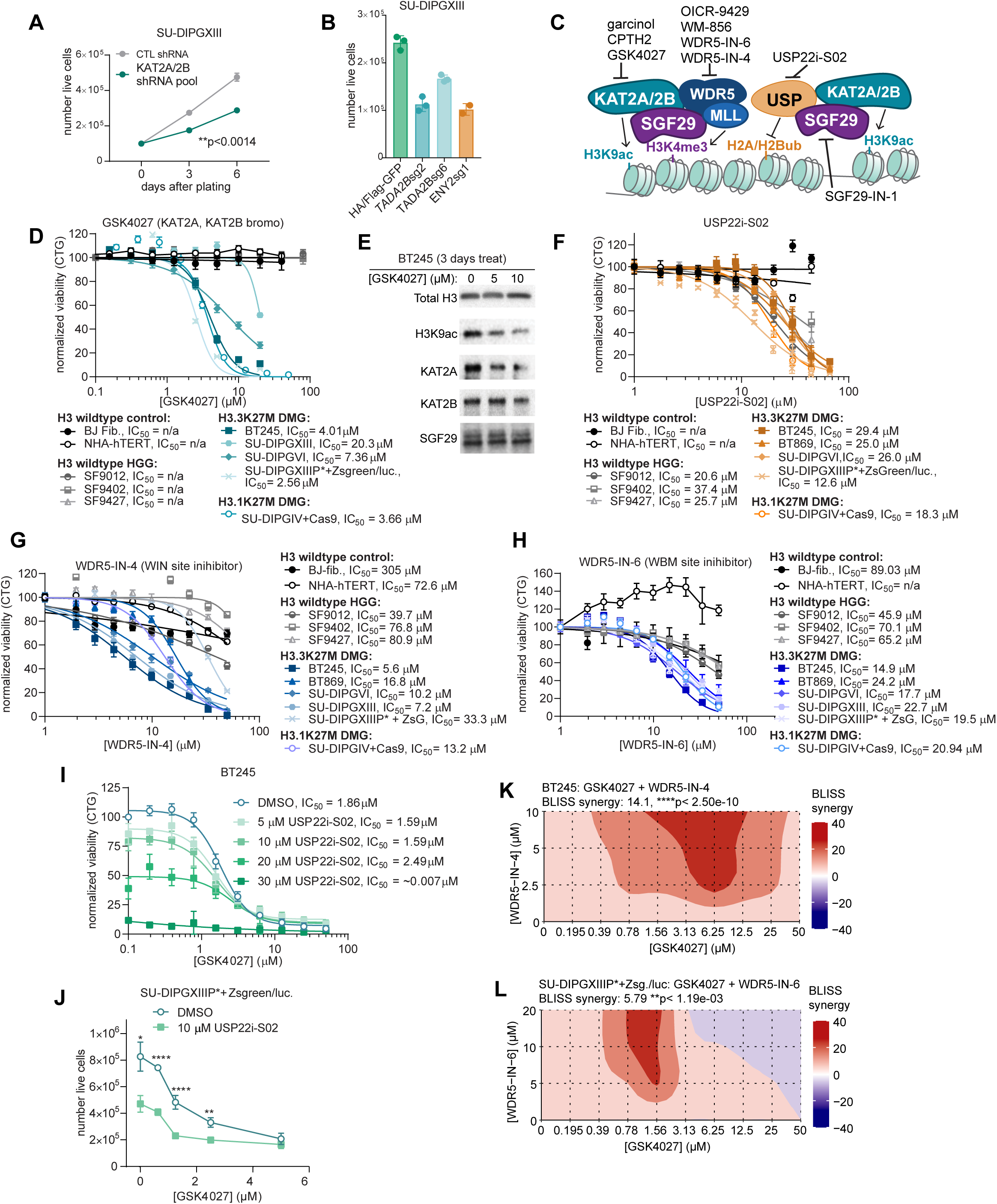
Genetic and pharmacological inhibition of SAGA/ATAC-associated chromatin modifying modules inhibits DMG growth. **A**, Growth curves of SU-DIPGXIII cells transduced with shRNAs to knockdown both *KAT2A* and *KAT2B,* showing significantly reduced growth compared to control scramble shRNA. **B**, Number of live SU-DIPGXIII+Cas9 cells expressing a control plasmid (HA/flag-GFP) or sgRNAs to KO SAGA-associated acetyltransferase activity (*TADA2B*), or de-ubiquitinase activity (*ENY2*). **C**, Schematic illustrating inhibition of histone-modifying modules of SAGA/ATAC complexes using chemical probes to target SAGA/ATAC-dependent histone acetylation (GSK4027, CPTH2, garcinol), SAGA-dependent H2A/H2B de-ubiquitination (USP22i-S02), or WDR5 inhibitors to target H3K4me3 methylation (OICR-9529, WM856, WDR5-IN-4, and WDR5-IN-6). **D**, Average CellTiter-Glo (CTG) cell viability following 7 days treatment with increasing doses of the KAT2A/2B bromodomain-targeting acetyltransferase inhibitor GSK4027 showing reduced growth of H3.3K27M and H3.1K27M mutant DMG cell viability (teal curves), but negligible effect on H3 wildtype HGG (grey curves) or non-transformed, H3 wildtype, control cells (black curves). **E**, Immunoblots confirming that GSK4027 treatment reduces H3K9ac without altering total H3 or SGF29 protein abundance in BT245 cells. **F**, Dose response curves for the DUB inhibitor USP22i-S02 showing selective inhibition of H3.3K27M and H3.1K27M mutant DMG cell viability relative to H3 wildtype pHGG and control cells. **G** and **H**, Dose response curves for the WDR5 WIN-site inhibitor WDR5-IN-4 (**G**) and the WDR5 WBM-site inhibitor WDR5-IN-6 (**H**), similarly showing selective DMG targeting. **I** and **J**, Average BT245 cell viability (**I**), and average number of live SU-DIPGXIIIP*+ZsG/luc cells (**J**) following treatment with combinations of GSK4027 and USP22i-S02 to simultaneously target both SAGA/ATAC-dependent histone acetyltransferase activity and SAGA-dependent H2A/H2B de-ubiquitination. **K** and **L**, BLISS synergy plots showing a combined effect of GSK4027 and WDR5-IN-4 treatment in reducing BT245 cell growth (\*\*\*\**p*<2.5×10^-10^, **K**) and of GSK4027 and WDR5-IN-6 treatment in reducing SU-DIPGXIIIP*+ZsG/luc growth (\*\**p*<1.74×10^-3^, **K**).

We also assessed roles for the SAGA-associated deubiquitinase (DUB) module involved in the removal of H2AK119ub and H2BK120ub (12,21,23) on DMG growth by generating *USP22KO* and *ENY2*KO cell lines. *USP22*KO alone does not significantly affect the growth of SU-DIPGVI cells (Supplementary Fig. S2F and S2G), potentially due to compensation by other DUBs like USP27x and USP51, as reported previously (23). However, *ENY2*KO, which is expected to simultaneously block USP22, USP27x, and USP51 recruitment to SAGA complexes (Fig. **1B**) (23,52), significantly inhibited DMG growth (Fig. **3B**). These results suggest that genetic targeting of the HAT and DUB modules of SAGA reduces DMG growth *in vitro*.

Tool compounds have been developed to target SAGA/ATAC-associated chromatin modifying enzymes, including members of the HAT, DUB, and H3K4me3-methylating modules (Fig. **3C**) (53–58). We find that the KAT2A/2B bromodomain inhibitor GSK4027 (59) selectively killed H3K27M mutant DMG cells compared to both non-transformed H3 wildtype control cells and H3 wildtype pHGG cells (Fig. **3D**). GSK4027 also dose-dependently reduced H3K9ac by immunoblotting, validating effective HAT inhibition (Fig. **3E**). In contrast, KAT2A/2B catalytic inhibitors (CPTH2 and garcinol) (54,60) impaired both DMG and non-transformed H3 wildtype cell growth (Supplementary Fig. S2H and S2I), potentially due to reported off-target effects on other acetyltransferases like p300 and CBP (54,60). Accordingly, we find that both H3K9ac and H3K27ac were reduced by CPTH2 treatment, suggesting broader inhibition of HAT activity (Supplementary Fig. S2J). GSK4027, garcinol, and CPTH2 treatment also reduced SU-DIPGXIII and BT245 live cell counts (Supplementary Fig. S2K and S2L), further validating a dose-dependent reduction in DMG growth due to KAT2A/2B inhibition using chemical probes with different mechanisms of action.

Treatment with USP22i-S02 to pharmacologically block SAGA-associated DUB activity (53) similarly inhibited the growth of H3.3K27M, H3.1K27M, and H3 wildtype pHGG cell growth with IC_50_ values in the ∼20 µM range, whereas non-malignant H3 wildtype pHGG were insensitive (Fig. **3F**). Finally, we determined DMG, H3 wildtype pHGG, and control cell sensitivity to WDR5 inhibitors designed to interface with either an arginine-binding cavity called the WIN site to block the recruitment of H3K4me3 methyltransferases (WDR5-IN-4) (56), or to bind to a hydrophobic cleft called the WBM site to impair MYC and N-MYC binding (WDR5-IN-6) (57). Both WDR5-IN-4 and WDR5-IN-6 selectively impaired DMG cell growth relative to both H3 wildtype controls and pHGG cell lines (Fig. **3G** and **H**). In contrast, alternative WDR5 inhibitors (OICR9429, WM586) (55,58) suppressed both H3.3K27M mutant DMG and control cell growth with similar IC_50_ values, indicating a lack of specificity for DMG (Supplementary Fig. S2M and S2N). In summary, these dose curve analyses reveal that DMG cells are sensitive to small molecules of SAGA/ATAC-associated chromatin modifying enzymes and point to GSK4027, SGF29-IN-1, WDR5-IN-4, and WDR5-IN-6 as selective inhibitors of DMG and/or pHGG growth (Supplementary Fig. S2O).

We then asked if there might be a combined effect of simultaneously inhibiting SAGA/ATAC-associated HAT-, DUB-, and WDR5-dependent histone modifying activities. Combined treatment with GSK4027 and USP22i-S02 to block both KAT2A/B and DUB chromatin modifying activities led to a more pronounced suppression of DMG cell growth than either inhibitor on its own (Fig. **3I** and **J**). Further drug combination studies reveal that co-treatment with GSK4027 and WDR5 inhibitors synergistically reduced DMG growth (Fig. **3K** and **L**), with greater synergy seen between GSK4027 and the WIN site-inhibitor WDR5-IN-4 (average BLISS score = 14.12). Overall, these results show that blocking SAGA/ATAC-dependent chromatin modifying enzymes selectively inhibits DMG growth, suggesting that these SAGA/ATAC-targeting chemical probes might be further developed for DMG treatment.

### SGF29 maintains active chromatin states in H3K27M mutant DMG

H3K27M oncohistones broadly reprogram the chromatin landscape of DMG, promoting hyperacetylation and other chromatin changes at genes that drive proliferation, survival, and stem/progenitor cell transcriptional programs (5,8,10,11,42,61,62). H3K27M oncohistones also restrict the propagation of H3K27me3 by impairing PRC2 catalytic efficiency and thereby blocking the lateral spread of repressive chromatin at certain target genes such as at tumor suppressor loci (4,5,63–65). To determine roles for SAGA and ATAC complexes in maintaining aberrant patterns of chromatin modifications in DMG, we applied tandem mass spectrometry to detect histone post-translational modifications (PTMs) on histones isolated by acid extraction from control and *SGF29*KO H3.3K27M DMG cell lines. These analyses reveal a reduction in histone H3 acetylation at lysines 9, 14, 18, and 23, associated with open chromatin, and decreased histone phosphorylation linked to mitotic progression (H3S10ph and H3S28ph) following *SGF29*KO in both HSJD-DIPG007 and SU-DIPGXIIIP* cells (Fig. **4A**; Supplementary Table 3) (66). To investigate the impact of SAGA/ATAC chromatin regulatory activity on DMG chromatin states genome-wide, we performed CUT&RUN in control and *SGF29*KO SU-DIPGXIII cells for SAGA/ATAC-associated H3K4me3 and H3K9ac and for H3K27M-altered H3K27me3. First focusing on histone modification changes associated with gene silencing, we performed differential peak calling, which identified 2,870 downregulated H3K9ac peaks (Fig. **4B**), 8,596 downregulated H3K4me3 peaks (Fig. **4C**), and 305 upregulated H3K27me3 peaks (Supplementary Fig. S3A) in the *SGF29*KO versus control SU-DIPGXIII cells. These data identify SAGA/ATAC complexes as key regulators of the active chromatin landscape established by H3K27M in DMG.

**Figure 4.**
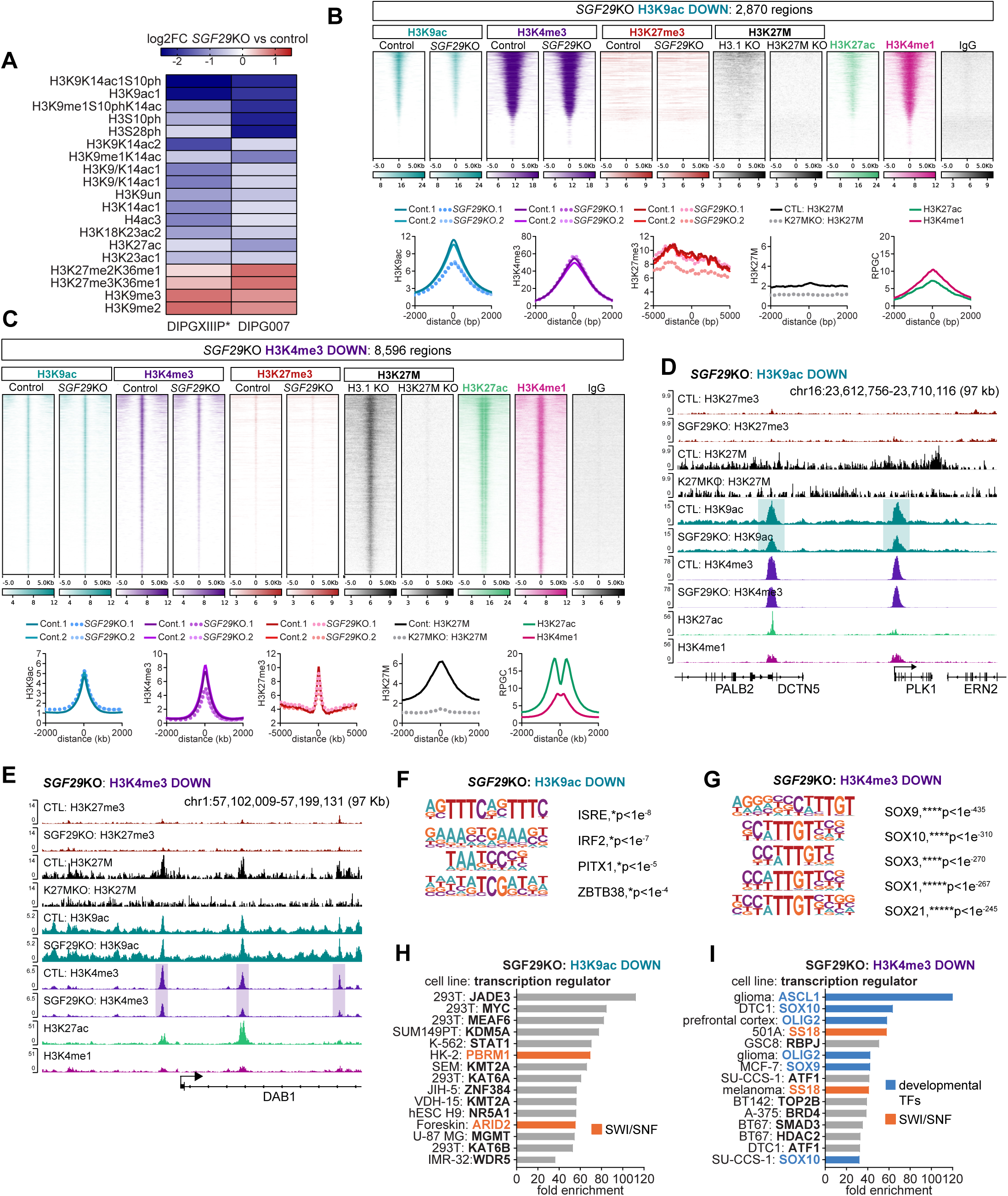
SGF29 sustains active chromatin states at both promoters and putative enhancers in H3K27M mutant DMG. **A**, Heatmap showing log_2_ fold change (log_2_FC) for various histone post-translational modifications in *SGF29*KO HSJD-DIPG007 and SU-DIPGXIIIP* cells compared to control cells as determined by mass spectrometry analysis of acid-extracted histones (see Methods). **B**, Heatmap (top panel) and average profile plots (bottom panel) of CUT&RUN tracks centered on significant H3K9ac-down peaks (2,870 regions, \**p*<0.01) in the *SGF29*KO SU-DIPGXIII cells. These H3K9ac-down peaks are enriched in H3K4me3, exhibit low H3K27M and H3K27me3 occupancy, and are marked by a poised enhancer-like state (H3K4me1-high, H3K27ac-low). **C**, Heatmap and profile plots of CUT&RUN tracks centered on the significant H3K4me3-down peaks (8,596 regions, \**p*<0.01), revealing H3K27M binding, low H3K27me3 enrichment and the presence of active enhancer marks (H3K4me1-high, H3K27ac-high). **D**, Genome browser snapshots of the *PLK1/DCTN5* (*Polo-Like Kinase 1*/*Dynactin 5)* loci showing representative H3K9ac-down peaks in the *SGF29*KO cells (teal boxes), which co-localize with both H3K4me3 and H3K4me1, but not H3K27M. **E**, Genome browser tracks showing H3K4me3-down peaks (purple boxes) in the promoter and gene body of the *DAB1* (*Disabled 1*) gene, which overlap with H3K27M, H3K27ac, and H3K4me1 peaks. In panels **B**-**E**, H3K27M CUT&RUN was performed in H3.1KO and H3K27MKO SU-DIPGXIII cells to control antibody specificity as previously described (63). **F** and **G**, Transcription factor binding sites identified using HOMER suggesting a significant enrichment of interferon-responsive transcription factor motifs (ISRE and IRF2) at the H3K9ac-down sites (**F**) and an enrichment in SOX family transcription factor motifs at the H3K4me3-down regions (**G**). **H** and **I**, Fold enrichment of known transcription factor binding sites from the ChIP ATLAS database within the H3K9ac-down (**H**) and H3K4me3-down (**I**) peak sets. This analysis suggests overlaps between SGF29-regulatated chromatin and binding sites for SWI/SNF chromatin remodeling complex binding (orange text and bars) and regions bound by developmental transcription factors like SOX9 and 10, ASCL1, and OLIG2 (blue text and bars).

To determine the relationship between H3K27M occupancy and SGF29-dependent chromatin regulation, we also performed H3K27M CUT&RUN in isogenic H3K27MKO and H3.1KO control SU-DIPGXIII cells, generated as previously described (63). Integrating the H3K27M CUT&RUN data with our H3K9ac, H3K4me3, and H3K27me3 profiling data reveals distinct chromatin fingerprints at SAGA/ATAC-regulated gene loci. We find that *SGF29*KO reduces H3K9ac at genes marked by constitutively high H3K4me3 and low H3K27M binding as illustrated by the chromatin profiles associated with the *PLK1/DCTN5* (*Polo-Like Kinase 1*/*Dynactin 5*; Fig. **4D**), *GPR139* (*G Protein Coupled Receptor 139*; Supplementary Fig. S3B), and the *BET1* (*Blocked Early in Transport 1*; Supplementary Fig. S3C) genes. In contrast, the H3K4me3-down peaks were marked by either H3K27M co-enrichment, as observed at the *DAB1* (*Disabled 1*) locus (Fig. **4E**), or by H3K4me3 and H3K27me3 co-enrichment as evident at the *NEUROD6* locus (*Neuronal Differentiation 6*; Supplementary Fig. S3D). Further analysis of these chromatin changes reveals a coordinated loss of gene-activating H3K9ac/H3K4me3 and gain of gene-silencing H3K27me3 at a subset of regions in the *SGF29*KO cells. As examples, we find that *SGF29* loss results in decreased H3K9ac and increased H3K27me3 at the *BET1* gene (Supplementary Fig. S3C), and both decreased H3K4me3 and increased H3K27me3 at the *NEUROD6* gene (Supplementary Fig. S3D).

We also determined whether these genomic regions that acquire a repressive chromatin state following *SGF29*KO might function as cis-regulatory promoters or enhancers. We first compared the H3K9ac-down, H3K4me3-down, and H3K27me3-up peak sets to previously generated H3K4me1 and H3K27ac enhancer mark ChIP-seq data from SU-DIPGXIII cells (6). The H3K9ac-down peaks exhibited a modest enrichment of histone PTMs found at poised enhancers (66) (H3K4me1-high and H3K27ac-low; Fig. **4B** and **D**, Supplementary Fig. S3B and S3C), whereas a subset of the H3K4me3-down peaks were associated with an active enhancer-like chromatin state (H3K27ac-high; Fig. **4C** and **E**). Consistent with these observations, genomic annotation reveals that both the H3K9ac-down and H3K4me3-down peaks localize not only to promoters and gene bodies but also to intergenic regions that may function as distal enhancers (Supplementary Fig. S3E). Lastly, in agreement with previous studies reporting a negligible overlap between H3K27ac- and H3K27me3-bound regions in DMG (40,63), the H3K27me3-up peaks showed negligible H3K27ac/H3K4me1 enrichment (Supplementary Fig. S3A). However, these *SGF29*KO-increased H3K27me3 peaks were frequently found in intergenic regions (Supplementary Fig. S3E), further implicating SGF29 in the regulation of distal enhancers.

To assess correlations between *SGF29*KO-mediated induction of repressive chromatin states and transcription factor activity in DMG, we performed motif enrichment analysis. We find that SGF29 maintains H3K9ac at regions containing interferon-sensitive response elements (ISREs) and consensus binding sites for IRF2 (Interferon Response Factor 2, Fig. **4F**). In contrast, the H3K4me3-down peaks overlapped with binding sites for SOX family transcription factors like SOX9 and SOX10, which regulate stem/progenitor cells and cell fate specification in the brain and other tissues (Fig. **4G**) (67). The H3K27me3-up peaks were similarly enriched in binding sites for developmental transcription factors like SOX14 and HOXC13 (Supplementary Fig. S3F).

Finally, comparing these regions to a database of published ChIP-seq results (ChIP ATLAS) (68), reveals an overlap between the H3K9ac- and H3K4me3-down peaks and binding sites for binding sites associated with the SWI/SNF ATP-dependent chromatin remodeling complexes (orange bars, Fig. **4H** and **I**), which have been linked to DMG stem cell function and malignancy (69). Consistent with the motif calling results (Fig. **4G**), the H3K4me3-down peaks were also enriched in binding sites for developmental transcription factors involved in glial and neuronal lineage specification, including ASCL1, OLIG2, and SOX family transcription factors (blue bars, Fig. **4I**).

### *SGF29*KO increases active chromatin marks and decreases gene-silencing H3K27me3 at a subset of regions

In addition to inducing chromatin states associated with gene repression, further analysis of our CUT&RUN profiling data suggests that *SGF29*KO also leads to gene-activating or transcriptionally permissive chromatin states at a subset of genomic regions. *SGF29*KO led to H3K9ac upregulation at 2,085 sites (Fig. **5A**), H3K4me3 upregulation at 861 sites (Fig. **5B**), and H3K27me3 downregulation at 2,224 regions (Fig**. 5C**). In contrast to the H3K9ac-down peaks, which were marked by low H3K27M binding (Fig. **4B**), the H3K9ac-up peaks showed moderate H3K27M occupancy and a robust enrichment in H3K4me1 and H3K27ac enhancer marks (Fig. **5A**). This pattern of co-localization between the H3K9ac-up peaks and H3K27M is evident at genes like *LAMB3 (Laminin Subunit β3)* and *LMNA* (*Lamin A/C*) (Fig. **5D** and Supplementary Fig. S4A). However, we observe minimal H3K27M binding at the upregulated H3K4me3 peaks, which instead exhibited a poised enhancer-like chromatin state (H3K4me1-high, H3K27ac-low; Fig. **5B**), as exemplified by *SGF29*KO-upregulated H3K4me3 peaks in the vicinity of a choline transporter *SLC44A3* (*Solute Carrier Family 44 Member 3*) (Fig. **5E**), a glutamate NMDA receptor (*GRIN3B*; Fig. **5F**), and the Dopamine D2 Receptor genes (*DRD2*; Supplementary Fig. S4B). Further examination of the *SLC44A3*, *GRIN3B*, and *DRD2* loci reveals a reduction in gene-silencing H3K27me3 accompanying the increased H3K4me3 (red and purple boxes; Fig. **5E** and **F**, Supplementary Fig. S4B), suggesting coordinated regulation of chromatin marks following *SGF29K*O at these loci.

**Figure 5.**
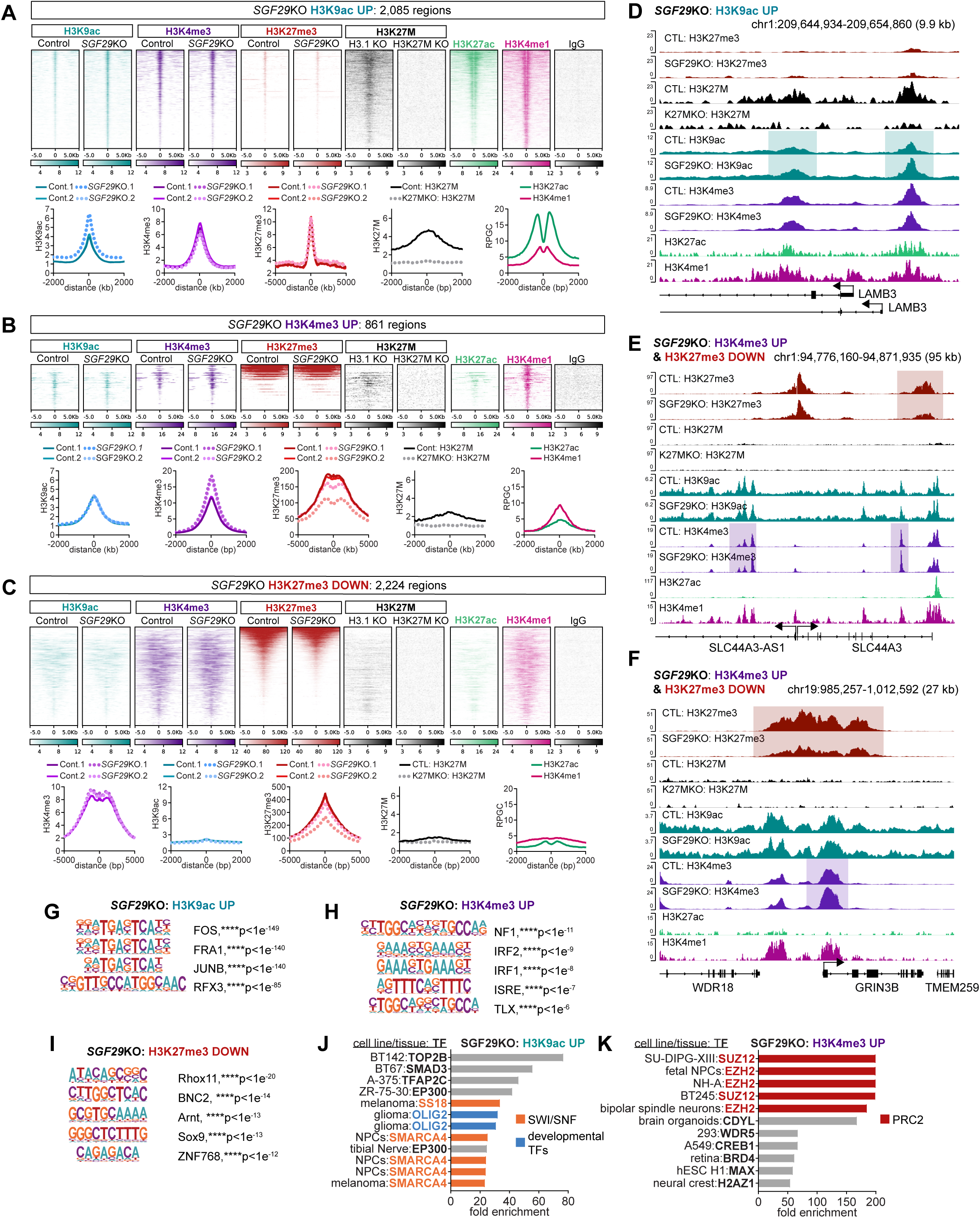
*SGF29*KO increases active chromatin marks and decreases gene-silencing H3K27me3 at a subset of cis-regulatory regions. **A**-**C**, Heatmap and average profile plots of CUT&RUN tracks from control versus *SGF29*KO SU-DIPGXIII cells showing significant H3K9ac-up peaks (2,085 regions, **A**), H3K4me3-up peaks (861 regions, **B**), and H3K27me3 up peaks (2,224 regions, **C**). Aligning these peak sets with H3K27M CUT&RUN data and H3K4me1/H3K27ac ChIP-seq data reveals distinct chromatin fingerprints: The H3K9ac-up peaks are enriched in H3K27M and exhibit an active enhancer-like state (H3K4me1-high, H3K27ac-high; **A**). The H3K4me3-up peaks show H3K27me3 co-occupancy and a poised enhancer state (H3K4me1-high, H3K27ac-low; **B**), whereas the H3K27me3-down peaks co-localized with H3K4me3, but lacked H3K4me1/H3K27ac enrichment (**C**). **D**-**F,** Genome browser snapshots illustrating genes exhibiting more active or transcriptionally permissive chromatin states in the *SGF29*KO cells. Panel **D** shows H3K9ac-up peaks (teal boxes) co-localizing with H3K27M and active chromatin marks at the *LAMB3* gene. Panels **E** and **F** show *SGF29*KO-increased H3K4me3 peaks (purple boxes) and decreased H3K27me3 peaks (red boxes) within the *SLC44A3* gene body (**E**) and at the *GRIN3B* promoter (**F**). **G** and **H,** Consensus transcription factor binding sites observed in the H3K9ac-up (**G**), H3K4me3-up (**H**), and H3K27me3-down (**I**) peaks from HOMER motif enrichment analysis demonstrating overlaps with AP-1, interferon-related, and developmental transcription factor binding sites. **J** and **K,** Fold enrichment of transcription factor and chromatin regulator binding sites, as catalogued in the ChIP ATLAS database, suggesting an overlap between the H3K9ac-up peaks and OLIG2 and SWI/SNF binding sites (**J**), and between the H3K4me3-up peaks and Polycomb Repressive Complex 2 (PRC2) binding sites (**K**).

We then examined the genomic distribution of regions that acquired more active chromatin features following *SGF29*KO. Consistent with our H3K4me1 and H3K27ac profiling, annotation of these regions reveals an association between the H3K9ac-up and H3K27me3-down peaks with distal, intergenic regions potentially acting as enhancers, and a strong overlap between the H3K4me3-up peaks and promoter regions (Supplementary Fig. S4C). Further motif enrichment analysis suggests a strong overlap between the H3K9ac-up peaks and binding sites for AP-1 family transcription factors like FOS and JUNB (Fig. **5G**), and an enrichment in IRF1/2 and ISRE binding sites among the H3K4me3-up peaks (Fig. **5H**), in line with our analysis of the H3K9ac-down peaks (Fig. **4F**). Lastly, the H3K27me3-down peaks were enriched in binding sites for developmental transcription factors like Sox9 and Rhox11 whose human orthologue, RHOXF1, regulates germ cell development (Fig. **5I**) (70). Finally, we find that *SGF29*KO induces H3K9ac at known binding sites for SWI/SNF and OLIG2 through comparisons with published ChIP-seq datasets (Fig. **5J**), whereas the H3K4me3-up and H3K27me3-down peaks were enriched in Polycomb Repressor Complex 2 (PRC2) binding sites (red text; Fig. **5K** and Supplementary Fig. S4D), suggesting dynamic remodeling of PRC2-dependent chromatin states. Taken together, these further chromatin profiling analyses demonstrate that *SGF29* loss not only leads to the repressive chromatin profiles described above (Fig. **4** and Supplementary Fig. S3) as anticipated, but also results in a more active chromatin state characterized by coordinated gains in H3K9ac and H3K4me3 and loss of H3K27me3 at a subset of genomic regions.

### SGF29 and KAT2A/2B acetyltransferases regulate the expression of transcripts associated with malignancy, metabolism, and brain development

To characterize the downstream transcriptional targets of SAGA/ATAC in DMG, we performed bulk RNA-seq on *SGF29*KO versus control SU-DIPGXIII and BT245 cells. These analyses identified differentially expressed genes (DEGs) across both cell lines using two independent sgRNAs (*SGF29*KO.A, *SGF29*KO.C) (Fig. **6A**, Supplementary Fig. S5A-S5C, Supplementary Table 4). In parallel, we generated RNA-seq data from SU-DIPGXIII cells treated for 25 hours with either vehicle (DMSO), the KAT2A/2B bromodomain inhibitor GSK4027, or the catalytic KAT2A/2B inhibitor CPTH2 (Fig. **6A**, Supplementary Fig. S5D-S5F, Supplementary Table 4). *SGF29*KO and HAT inhibitor (HATi) treatment led to both statistically significant up- and downregulated genes with a partial overlap between the DEGs observed following *SGF29*KO or HATi treatment (Fig. **6A**; Supplementary Fig. S5A-S5H). These transcriptional changes are consistent with our chromatin profiling data identifying numerous genomic regions marked by either repressive (Fig. **4**; Supplementary Fig. S3) or gene-activating (Fig. **5**, Supplementary Fig. S4) chromatin alterations in the *SGF29*KO versus control SU-DIPGXIII cells. Comparing these DEGs to the chromatin profiling data reveals that the transcription start sites (TSS) of a subset of the silenced genes were localized near (+/-30 kb) the H3K9ac- and H3K4me3-down peaks from the CUT&RUN chromatin profiling, but minimal overlap with the H3K27me3-up peaks (Fig. **6B**, Supplementary Fig. S5I). In contrast, a subset of the upregulated genes were marked by a loss of H3K27me3 and/or increased H3K9ac and H3K4me3 (Fig. **6C**, Supplementary Fig. S5J). Together, these integrative analyses demonstrate a partial overlap between SGF29-dependent chromatin and transcriptional changes in DMG.

**Figure 6.**
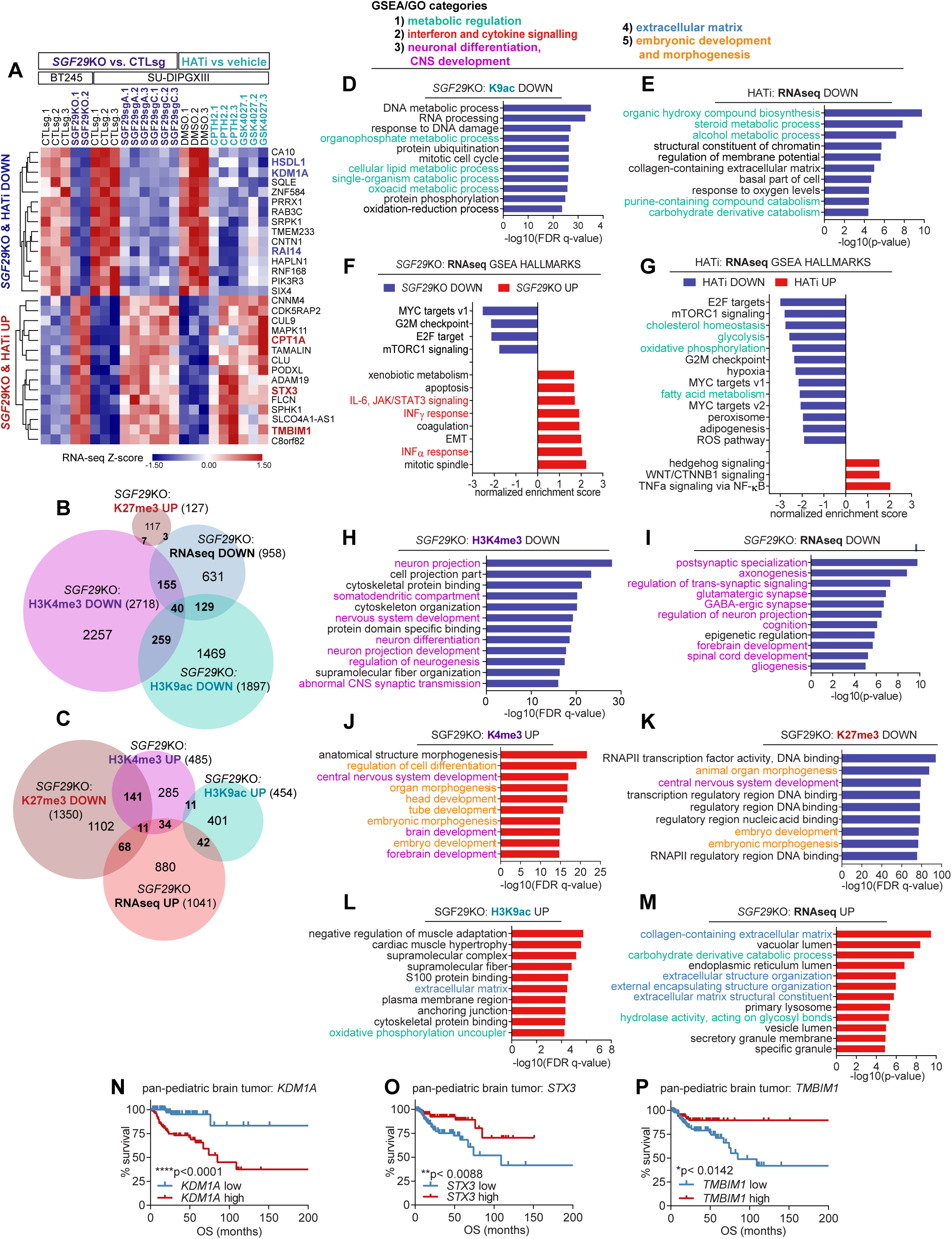
SGF29 and KAT2A/2B regulate the expression of transcripts associated with malignancy, metabolism, and development. **A,** Heatmap of RNA-seq Z-scores summarizing differentially expressed genes (DEGs) in *SGF29*KO SU-DIPGXIII and BT245 DMG cells and in SU-DIPGXIII cells following vehicle (DMSO) or KAT2A/2B inhibitor treatment (GSK4027 and CPTH2). **B,** Venn diagram illustrating the overlap between the significantly downregulated transcripts from the RNA-seq analysis of *SGF29*KO versus control cells and the H3K9ac-down, H3K4me3-down, and H3K27me3-up peaks (see Fig. 4B and **C**, Supplementary Fig. 3A). **C,** Venn diagram showing partial overlap between the upregulated transcripts from the RNA-seq analysis and the H3K9ac-up, H3K4me3-up, and H3K27me3-down peaks from the CUT&RUN profiling (see Fig. 5A-**C**). **D** and **E,** GO analysis of genes located within ±30 kb of the H3K9ac-down peaks (**D**) and associated with the HATi-downregulated transcripts (**E**). **F** and **G,** Pre-ranked GSEA of the RNA-seq analysis of the *SGF29*KO versus control cells (**F**) and in the HATi-treated versus vehicle-treated condition (**G**), revealing a downregulation of MYC and mTOR target genes and metabolic genes associated with malignancy. **H** and **I,** GO analysis of genes located within ±30 kb of the H3K4me3-down peaks (**H**), or showing decreased expression upon *SGF29*KO in the RNA-seq analysis (**I**). **J**-**L,** GO analysis of genes located within ±30 kb of H3K4me3-up peaks (**J**), the H3K27me3-down peaks (**K**), or the H3K9ac-up peaks (**L**), or showing increased expression upon *SGF29*KO in the RNA-seq analysis (**M**). **N**-**P**, Kaplan-Meier survival curves grouping pediatric brain tumors by high and low protein expression of SAGA/ATAC gene targets suggesting that increased expression of the SAGA/ATAC-induced gene, *KDM1A,* correlates with worse clinical outcomes (**N**), whereas the SAGA/ATAC-repressed genes, *STX3* and *TMBIM1,* correlate with improved survival (**O** and **P**).

We applied Gene Ontology (GO) and Gene Set Enrichment Analysis (GSEA) to identify overrepresented gene categories near *SGF29*KO-altered H3K4me3, H3K9ac, and H3K27me3 peaks and amongst the DEGs identified by RNA-seq (Fig. **6D**-**M**). GO analysis reveals that *SGF29*KO results in a loss of H3K9ac near genes regulating mitotic progression, DNA replication, and metabolism (Fig. **6D** and **E**). Consistently, *SGF29*KO and HATi treatment reduced the expression of metabolic genes, G2/M checkpoint markers, and oncogene-induced transcriptional signatures downstream of MYC, E2F, and mTORC1 (Fig. **6F** and **G**; Supplementary Fig. S5K and S5L). These transcriptional changes are reminiscent of previous studies, similarly identifying genes associated with cell cycle progression and oncogenic signaling/transcription as downstream targets of SAGA/ATAC in other cancers (13,71–74). *SGF29*KO also upregulated genes involved in interferon signaling (Fig. **6F**), consistent with our motif calling analyses identifying IRF2 and ISRE binding sites among the H3K9ac-down and H3K4me3-up peaks (Fig. **4F** and Fig. **5H**). Additional analyses suggest that *SGF29*KO reduces H3K4me3 at neuronal differentiation and synaptogenesis genes and correspondingly represses the expression of transcripts related to brain development and cognition (Fig. **6H** and **I**, Supplementary Fig. S5M and S5N). The H3K4me3-up and H3K27me3-down peaks were similarly associated with CNS and embryonic development (Fig. **6J** and **K**), further supporting a role for SGF29 in coordinating chromatin states at developmental genes. Finally, we find that extracellular matrix (ECM) genes were found near the *SGF29*KO-induced H3K9ac peaks and correspondingly upregulated in the RNA-seq analysis (Fig. **6L** and **M**).

Additional analyses suggest that SGF29 and KAT2A/2B promote the transcription of genes that are overexpressed in pediatric HGG versus LGG patient tumor samples, like *RAI14* (*Retinoic Acid Induced 14*), *HSDL1* (*Hydroxysteroid Dehydrogenase Like 1*), and *KDM1A* (*Lysine Specific Demethylase 1*; Supplementary Fig. S5O), as these genes are downregulated by both *SGF29*KO and HATi treatment (bold blue text, Fig. **6A**). In contrast, transcripts upregulated by *SGF29*KO and HATi treatment were instead overexpressed in LGG compared to HGG, including *CPT1A* (*Carnitine Palmitoyltransferase 1A*), *STX3* (*Syntaxin 3*), and *TMBIM1* (*Transmembrane BAX Inhibitor Motif Containing 1*, bold red text; Fig. **6A**, Supplementary Fig. S5P). Finally, Kaplan-Meier survival analyses show a significant correlation between expression of the SAGA/ATAC target genes (*KDM1A*, *STX3*, and *TMBIM1*) and disease outcomes in a pan-pediatric brain tumor proteomics dataset (Fig. **6N**-**P**). Together, these results indicate that SGF29 and SAGA/ATAC-associated acetyltransferases regulate the transcription of DMG disease-associated genes governing cell cycle, metabolism, and both CNS and embryonic development.

### Combinations of KAT2A/2B and cholesterol synthesis inhibitors synergistically reduce DMG growth

We next sought to determine if selected SAGA/ATAC target genes from our RNA-seq analysis might mediate the growth defects observed due to SAGA/ATAC inhibition. Our chromatin and transcriptional profiling data point to SAGA/ATAC-dependent regulation of key genes involved in promoting cell growth, including metabolic regulators like the TCA cycle isocitrate dehydrogenase genes, *IDH1* and *IDH3A,* which showed reduced H3K9ac binding upon *SGF29*KO (Supplementary Fig. S6A and S6B). *SGF29*KO similarly led to a loss of H3K9ac at gene loci encoding mevalonate pathway genes involved in cholesterol synthesis and steroidogenesis like *SQLE* (*Squalene Epoxidase*) and *HMGCS1* (*HMG-CoA Synthase 1*; Supplementary Fig. S6C and S6D). Consistently, both *SGF29*KO and HATi treatment led to a downregulation of mRNAs encoding rate-limiting enzymes necessary for cholesterol synthesis (Fig. **7A** and **B**), which was previously identified as a metabolic dependency in DMG and other brain tumors (47,75). SAGA/ATAC-regulated cholesterol synthesis genes include rate-limiting enzymes like *FDFT1* (*Farnesyl-diphosphate Farnesyltransferase 1*), *SQLE*, and *LSS* (*Lanosterol Synthase;* Fig. **7A** and **B**). These results further highlight metabolic genes as downstream targets of SAGA/ATAC-dependent chromatin regulation in DMG.

**Figure 7.**
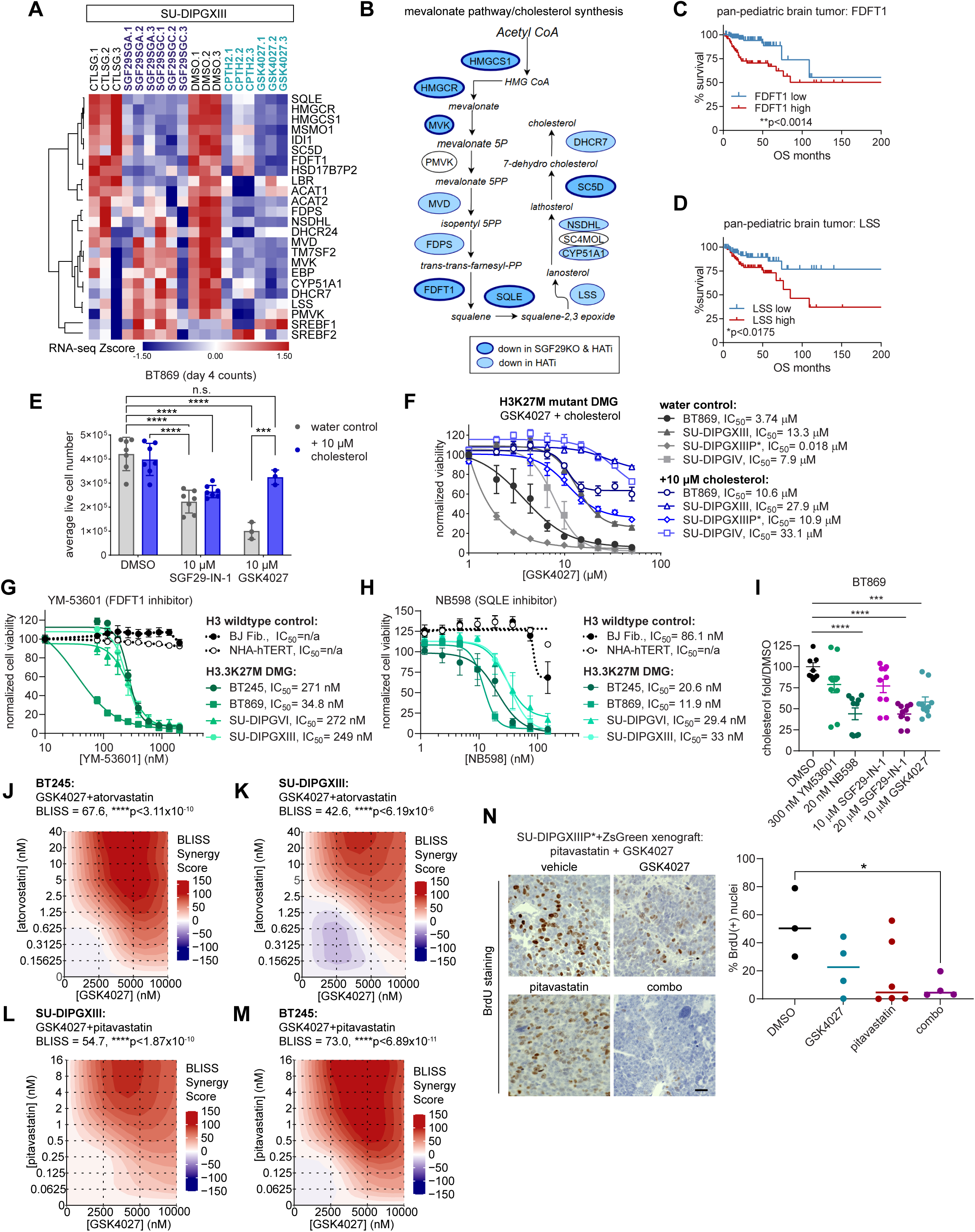
Combinations of KAT2A/2B inhibitors and cholesterol synthesis inhibitors synergistically reduce DMG growth. **A,** Heatmap summarizing RNA-seq Z-scores of mevalonate pathway and cholesterol metabolism genes, which were downregulated by pharmacological inhibition of KAT2A/2B and by *SGF29*KO in DMG cells. **B,** Pathway diagram of cholesterol synthesis showing genes that were downregulated by HATi (shaded blue) or by both HATi and *SGF29*KO (outlined in blue). **C** and **D,** Kaplan-Meier survival curves from pan-pediatric brain tumor patient proteomics data showing that elevated protein levels of the cholesterol synthesis enzymes farnesyl-diphosphate farnesyltransferase 1 (FDFT1; **C**), and lanosterol synthase (LSS; **D**), correlate with poor clinical outcomes. **E,** Live BT869 DMG cell counts after 4 days treatment with SGF29-IN-1 and GSK4027 either alone (water control) or supplemented with 10 µM water soluble cholesterol (cholesterol-methyl-β-cyclodextrin), indicating that cholesterol add-back can rescue KAT2A/2B inhibition, but not SGF29 inhibition. **F**, CellTiter-Glo dose curves from H3.3K27M and H3.1K27M mutant DMG cells treated with increasing doses of GSK4027 ± cholesterol, revealing decreased sensitivity to GSK4027 with cholesterol supplementation. **G** and **H,** CellTiter-Glo dose curves showing average DMG (green curves) and control cell (black curves) viability following 7 days of treatment with the FDFT1 inhibitor YM-53601 (**G**), or with the squalene epoxidase (SQLE) inhibitor NB598 (**H**). **I,** Average fold change in total cholesterol in BT869 DMG cells treated with inhibitors of cholesterol synthesis or small molecules targeting SGF29- and KAT2A/2B-dependent chromatin regulation, as determined using an AMPLEX-red fluorescent cholesterol assay and normalized to vehicle control (see Methods). **J**-**M**, Heatmaps of BLISS synergy scores from dose curve matrices of GSK4027 and either the HMGCR inhibitor atorvastatin (**J** and **K**), or pitavastatin (**L** and **M**) to simultaneously inhibit KAT2A/2B activity and cholesterol synthesis in BT245 and SU-DIPGXIII cells. **N** and **O**, Representative images (left panel) and quantification (right panel) of BrdU-labelled S-phase nuclei in SU-DIPGXIIIP*+ZsG/luc orthotopic xenografts following treatment with 10 mg/kg pitavastatin, 10 mg/kg GSK4027, or a combination for 6 weeks (\**p*<0.05, ANOVA). Scale bar is 25 microns.

To explore whether genes regulating the mevalonate pathway and cholesterol metabolism might represent unique DMG-specific vulnerabilities, we similarly analyzed pan-cancer CRISPR dependency screen data from DepMap. Genes like *HMGCS1* and *HMGCR* (*3-Hydroxy-3-Methylglutaryl-CoA Reductase*) were revealed as strong cancer dependencies regardless of the tissue of origin, whereas loss of *FDFT1* and *SQLE* had moderate effects on cancer cell growth overall (Supplementary Fig. S7A). Additional analysis of the pan-pediatric brain tumor patient proteomics data reveals that proteins linked to cholesterol metabolism are differentially expressed in HGG versus LGG (Supplementary Fig. S7B) and show a correlation between elevated FDFT1 and LSS protein and reduced patient survival (Fig. **7C** and **D**). These findings suggest that cholesterol biosynthesis enzymes represent metabolic dependencies associated with malignancy in both adult cancers and pediatric glioma, consistent with previous findings from studies in DMG and other cancers (41–44,46).

We then asked whether downregulation of cholesterol biosynthesis genes contributes to the growth defects induced by SAGA/ATAC inhibition. We performed rescue experiments by supplementing the DMG growth media with water-soluble cholesterol as previously described (43). While cholesterol supplementation was not sufficient to overcome growth repression induced by SGF29-IN-1, which broadly inhibits SAGA/ATAC activity, we find that cholesterol add-back rescued GSK4027-mediated DMG growth inhibition (Fig. **7E** and **F**). These results support a role for cholesterol metabolism downstream of SAGA/ATAC-dependent gene regulation via KAT2A/2B acetyltransferase activity.

To assess the impact of KAT2A/2B-dependent regulation of cholesterol metabolism genes on DMG growth, we then treated DMG cells with various cholesterol pathway-targeting drugs either alone or in combination with GSK4027, chosen as a SAGA/ATAC inhibitor based on strong selectivity for DMG (Supplementary Fig. S2O). Treatment with an FDFT1 inhibitor (YM-53601) (76) or with two structurally distinct SQLE inhibitors (NB598 and terbinafine) (77) for 7 days selectively repressed DMG compared to control cell growth (Fig **7G** and **H**, Supplementary Fig. 7C). A 24-hour treatment with the SQLE inhibitor NB598 (positive control), or with SAGA/ATAC-targeting GSK4027 or SGF29-IN-1 also resulted in a significant reduction in total cholesterol produced by BT869 cells (Fig. **7I**), confirming that these small molecules effectively suppress cholesterol synthesis. Additional studies show that DMG cells are also sensitive to statins targeting HMGCR activity like atorvastatin and pitavastatin (Supplementary Fig. S7D and S7E), confirming a previous study (41). However, both H3K27M mutant DMG and non-transformed control cells exhibited similar sensitivities to statins (Supplementary Fig. S7D and S7E). Together, these results point to cholesterol metabolism as a key driver of DMG growth acting downstream of KAT2A/2B-dependent regulation of histone acetylation and transcription.

Given that DMG cells were more sensitive to GSK4027 treatment than control cell lines (Fig. **3D**, Supplementary Fig. S2O) and our observation that GSK4027 treatment strongly inhibited the transcription of cholesterol and steroid metabolism genes (Fig. **6E**, Fig. **7A** and **B**), we reasoned that GSK4027 would sensitize DMG cells to cholesterol synthesis inhibitors by downregulating key metabolic regulators. Indeed, treatment with combinations of GSK4027 and either atorvastatin or pitavastatin resulted in a synergistic reduction in SU-DIPGXIIIP*, SU-DIPGXIII, and BT245 growth (Fig. **7J-M**, Figure S7F and S7G). Following up on previous studies showing that brain penetrant statins can reduce glioma growth both *in vitro* and *in vivo* (41), we then asked whether combined inhibition of SAGA/ATAC and cholesterol biosynthesis can suppress DMG growth in a xenograft mouse model. SU-DIPGXIIIP* xenograft-bearing NSG mice were treated by IP injection for 6 weeks with vehicle, 10 mg/kg pitavastatin, 10 mg/kg GSK4027, or a combination, followed by BrdU labeling to assess tumor cell proliferation. Immunohistochemical analysis revealed a significant reduction in BrdU-positive cells in DMG tumors treated with the combination of pitavastatin and GSK4027 relative to controls, whereas individual compounds did not significantly reduce BrdU labelling (Fig. **7N**). Taken together, these studies demonstrate that concurrent targeting of SAGA/ATAC function and cholesterol biosynthesis cooperatively suppresses DMG growth both *in vitro* and *in vivo*.

## Discussion

H3K27M mutant histone incorporation into chromatin is associated with increased histone acetylation and altered H3K4me3 associated with active transcription (2,6,7,9,42,62), but the enzymes and chromatin regulators required for this hyperactive chromatin state have remained elusive. We identified the SAGA/ATAC-associated chromatin reader, SGF29, as a key driver of H3K27M mutant DMG cell growth and tumorigenesis. Our further studies show that small molecule inhibitors of SAGA/ATAC-associated chromatin modifying modules involved in histone H3 acetylation, H2A/H2B de-ubiquitination, and the regulation of H3K4me3 via WDR5 similarly reduce DMG growth. Many of the SAGA/ATAC-targeting small molecules selectively inhibited DMG growth compared to control cell lines, raising the possibility that these tool compounds targeting SAGA/ATAC activity might be further developed as selective DMG therapeutics. Collectively, our genetic and pharmacological targeting studies establish SGF29 and SAGA/ATAC-associated chromatin-modifying modules as actionable chromatin dependencies in DMG and highlight their potential as targets for therapeutic intervention.

The coordinated activity of SAGA/ATAC complexes is essential for maintaining cellular differentiation states in normal development and cancer (13,15,27,35,51,78). Integrative analysis of RNA-seq and chromatin profiling from SGF29-deficient cells demonstrates that SGF29 orchestrates chromatin states at genes governing cancer metabolism, malignancy and both embryonic and neurodevelopmental programs in DMG, consistent with studies identifying TRRAP and USP22 as regulators of brain tumor initiating cell function and gliomagenesis (27,34,79) and linking SAGA/ATAC-dependent gene regulation to cancer metabolism (80–83). Specifically, SGF29 loss results in reduced H3K9ac at genes involved in mitotic progression, DNA replication, and metabolism, while H3K4me3 and H3K27me3 changes frequently occur at loci associated with brain development and embryogenesis. Further GSEA of SGF29- and KAT2A/2B-regulated genes identifies MYC- and mTOR-associated transcriptional programs as downstream targets of SAGA/ATAC in DMG, aligning with previous studies implicating these complexes in oncogenic transcriptional networks and growth-promoting signaling pathways in multiple myeloma, neuroblastoma and other cancers (14,30,32,35,71,72). Together, these findings establish SGF29-dependent SAGA/ATAC activity as a key determinant of the DMG chromatin landscape that modulates distinct histone modifications at metabolic genes (H3K9ac) and at embryonic/neurodevelopmental genes (shifts in H3K4me3 and H3K27me3 enrichment).

Comparing our CUT&RUN results and genome-wide H3K27M binding data reveals that a subset of regions exhibiting *SGF29*KO-dependent chromatin changes overlap with H3K27M-bound regions, including the H3K4me3-down peaks and a subset of the differential H3K9ac peaks. These data are correlative, but suggest an association between SAGA/ATAC-dependent chromatin regulation and H3K27M binding at a subset of *cis* regulatory elements. Consistently, we find that H3K27M mutant DMG cells are more sensitive to pharmacological SAGA/ATAC-targeting than either H3 wildtype control or pHGG tumor cells, further implicating SAGA/ATAC in H3K27M mutant cell growth. However, a direct link or causal relationship between H3K27M and SAGA/ATAC activity has not yet been established. Additional studies are needed to determine the extent to which SAGA/ATAC-dependent chromatin and transcriptional regulation influences H3K27M chromatin occupancy or, conversely, whether chromatin alterations induced by H3K27M oncohistone incorporation facilitate SAGA/ATAC recruitment and chromatin regulatory activity in DMG.

Another limitation of the current study is that direct binding sites for SGF29 and other core SAGA/ATAC chromatin regulators have not yet been identified in H3K27M mutant DMG, making it difficult to distinguish between the direct and indirect effects of SGF29 loss on DMG chromatin states. For example, *SGF29*KO induced both chromatin changes at binding sites for numerous developmental transcription factors, including SOX9, SOX10, and OLIG2, and altered the expression of developmental regulators by RNA-seq. It is possible that some of the chromatin-state and metabolic vulnerabilities explored in this study may be secondary to altered expression of developmental transcription factors and DMG cell fate changes, such as shifts in H3K4me3/H3K27me3 bivalent domains toward either more active or repressed chromatin states, or cell-state dependent changes in metabolic demand as previously reported (41).

Altered metabolism is a hallmark of brain cancer, with recent studies reporting aberrant mitochondrial activity, disrupted glucose and glutamine uptake and metabolism, and dysregulated cholesterol metabolism in DMG (41–44,47). In yeast, SAGA associates with ergosterol biosynthesis genes under low-sterol conditions and subsequently recruits the SWI/SNF chromatin remodeling complex to regulate the expression of sterol biosynthesis enzymes (84) and KAT2A has recently been linked to *FDFT1* regulation in liver cancer (80). Several recent studies have identified cholesterol metabolism as a key metabolic vulnerability in H3K27M mutant DMG (41,43,44). However, little is known about the chromatin and transcriptional regulators that sustain the expression of metabolic enzymes supporting DMG growth. We show that chemical inhibition of KAT2A/2B and SGF29 knockout alter the expression of cholesterol biosynthesis genes, and cholesterol add-back rescue experiments indicate that these metabolic defects arise downstream of KAT2A/2B-dependent histone acetylation. Additional studies demonstrate that DMG cells are highly sensitive to brain-penetrant cholesterol-targeting therapies administered either alone or in combination with GSK4027, which blocks SAGA/ATAC-dependent KAT2A/2B chromatin binding through their bromodomains. Together, these findings provide a rationale for further development of therapeutic approaches that target SAGA/ATAC-dependent chromatin regulation, both alone and in combination therapies, for the treatment of H3K27M mutant DMG.

## Supporting information

Supplementary Data Table 4: RNA-seq summary

Supplementary Data Table 3: Histone mass spectrometry analysis

Supplementary Data Table 2: CRISPR screen summary

Supplementary Data Table 1: sgRNA and shRNA sequences

## Data Availability

Sequencing data generated in this study is available on the Gene Expression Omnibus (GEO) under accession numbers GSE317275, GSE317235 and GSE318286. H3K4me1 and H3K27ac ChIP-seq data used in this study are available on GEO under accession number GSE110572.

## Authors’ Contributions

**R. Richard:** Methodology, investigation, validation, data curation, analysis, writing-review and editing. **C. Bagnetto:** Investigation, validation, data curation, visualization, analysis, writing-review and editing. **R. Murdaugh.** Methodology, investigation, validation, analysis. **B. Eberl:** Methodology, investigation, validation, analysis. **A. Jiao:** Investigation, validation, analysis, writing-review. **A. Kebede:** Investigation, validation, analysis, writing-review. **E. Campos-Hensley:** Investigation, visualization. **B. Zee:** Investigation, validation, analysis, writing-review. **M. Harrington:** Investigation, validation, analysis. **M. Filbin:** Conceptualization, resources. **A. Serin-Harmanci:** Methodology, formal analysis. **Y. Shi:** Conceptualization, resources, writing-review and editing. **J. Anastas:** Conceptualization, resources, data curation, formal analysis, supervision, funding acquisition, validation, visualization, methodology, writing-original draft, project administration, writing-review and editing.

## Acknowledgements

These studies were supported by funding from the National Institutes of Health (5R01NS129860), the Cure Starts Now Foundation, and the Rally Foundation for Pediatric Cancer Research and Kids Join the Fight, the Caroline Weiss Law Fund for Research in Molecular Medicine, and the ChadTough Defeat DIPG Foundation. Dr. Rebecca Murdaugh and Caitlin Bagnetto received training support from the Baylor College of Medicine (BCM) Center for Cell and Gene Therapy (CAGT) training grant (NHLBH, 5T32HL092332). Caitlin Bagnetto received additional support from the BCM Training Program in Cell and Molecular Biology (NIGMS, T32GM136560). We thank Dr. Bruno DiStefano for the TADA2B and ENY2 sgRNA plasmids. Histone mass spectrometry was performed at Taplin Mass Spectrometry Facility at Harvard Medical School. We thank Dr. Daniel Kraushaar and Dr. Kieu Pham from the Genomic and RNA Profiling Core at Baylor College of Medicine for their support in conducting CUT&RUN and RNA-seq studies with the support of funding from the NIH NCI (P30CA125123) and CPRIT (RP250580) grants as well as the sequencing core at Tufts University. We also thank Dr. Christopher Ward and the Mouse Metabolism and Phenotyping Core (NIH UM1HG006348, NIH R01DK114356, and NIH R01HL130249) and the Smith Breast Center Pathology lab for their assistance in live animal imaging and histopathological studies. This project was also supported by the Cytometry and Cell Sorting Core at Baylor College of Medicine with funding from the CPRIT Core Facility Support Award (CPRIT-RP240432) and the NIH (CA125123, OD036336, and OD038251), and the assistance of Joel M. Sederstrom.

---

**Supplementary Figure 1.**
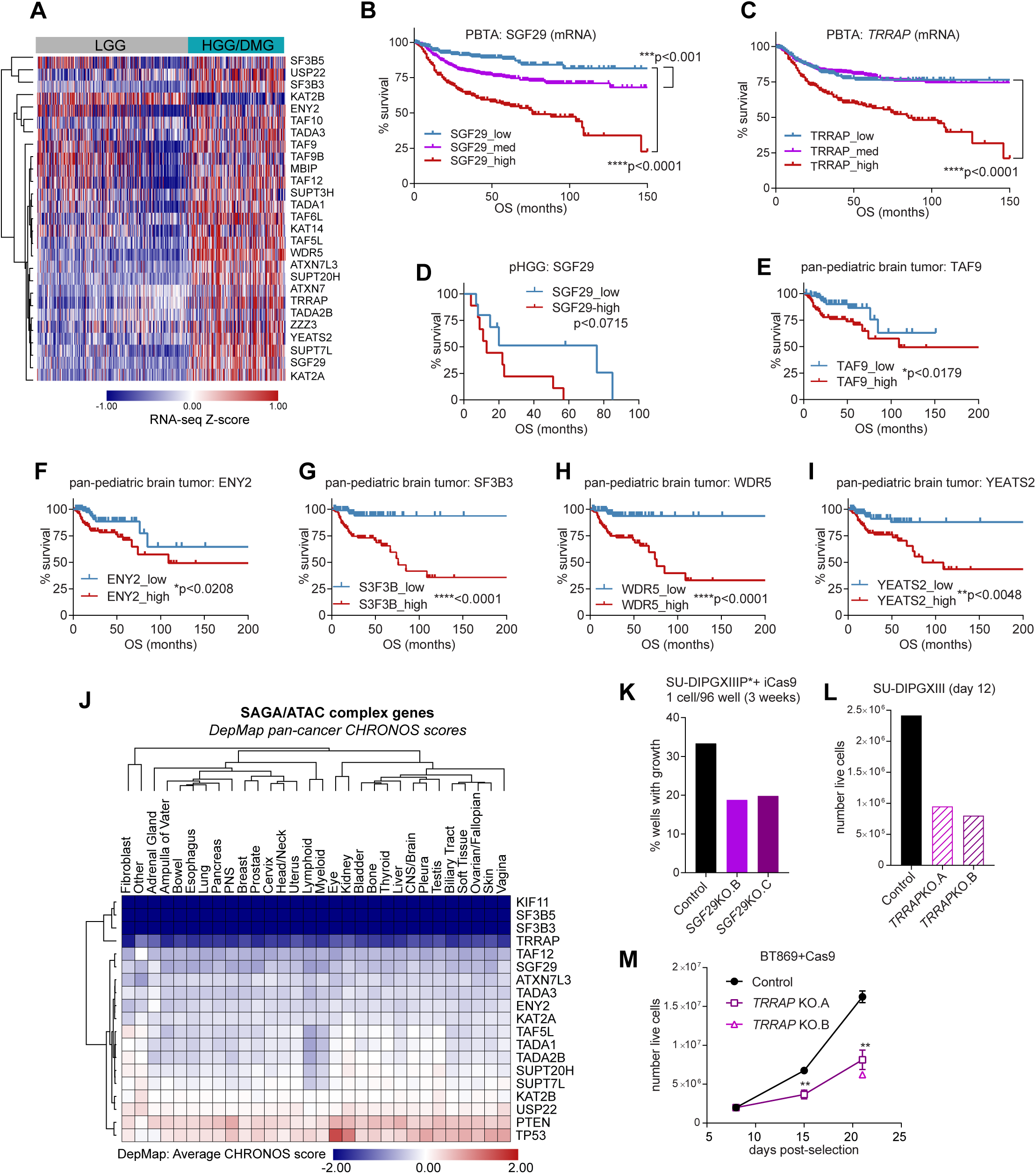
Increased expression of SAGA/ATAC genes is associated with poor clinical outcomes in pediatric brain tumors**. A,** Heatmap of Z-scores from publicly available RNA-seq analysis of primary tumors/diagnostic patient samples in the pediatric brain tumor atlas (PBTA), suggesting increased SAGA/ATAC-related mRNA expression in high grade glioma (HGG) versus low grade glioma (LGG). **B** and **C**, Kaplan-Meier survival curves grouped by low, medium, and high *SGF29* (**B**) and *TRRAP* (**C**) mRNA levels in patient tumors. **D**-**I**, Kaplan-Meier survival analyses of pediatric brain tumor patients grouped according to low and high SAGA/ATAC primary tumor protein expression from CPTAC (diagrammed in Fig. 1B). Increased expression of SAGA-associated TAF9, ENY2 and SF3B3 was associated with decreased survival (**E**-**G**) as were elevated levels of ATAC-associated WDR5 and YEATS2 (**H** and **I**). **J**, Heatmap of average CHRONOS scores from DepMap pan-cancer CRISPR screen data showing genetic dependencies on SAGA/ATAC genes as compared to universal dependency (*KIF11*, positive control) and *TP53* and *PTEN* included as example tumor suppressors. **K**, Clonal expansion of single SU-DIPGXIIIP*+iCas9 cells grown from single cells in 96-wells, indicating decreased growth and expansion of *SGF29*KO clones. **L**, Number of live SU-DIPGXIII+Cas9 cells 12 days after starting selection to knockout *TRRAP* with two independent sgRNAs. **M**, Growth curves of control and *TRRAP*KO BT869+Cas9 cells, indicating significantly reduced growth of the KO cells.

**Supplementary Figure 2.**
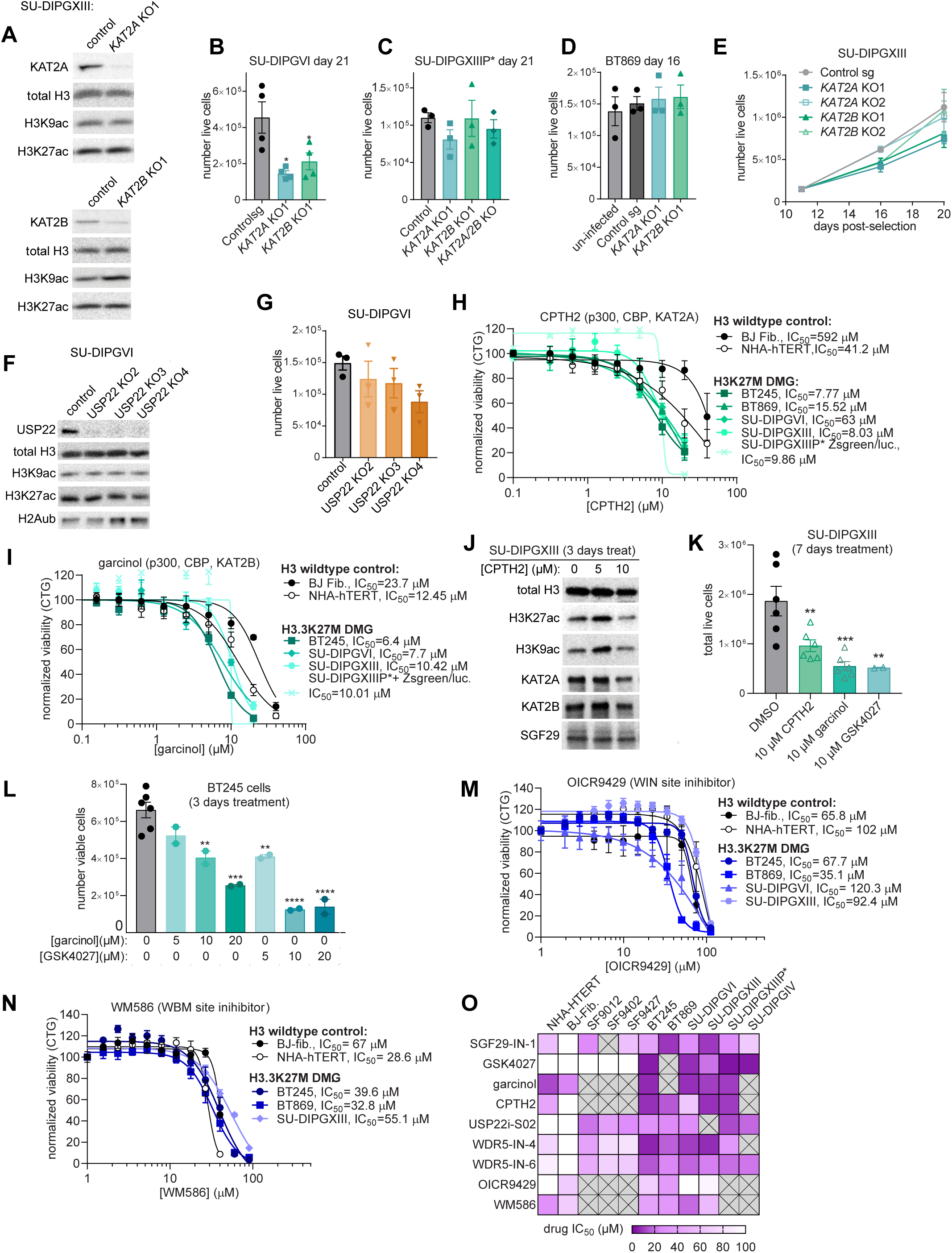
Inhibition of SAGA/ATAC-associated chromatin modifying modules reduces DMG growth. **A**, Immunoblot analysis of whole cell lysates from control, *KAT2A*KO (top panel), or *KAT2B*KO (bottom panel) SU-DIPGXIII+Cas9 cells showing reduced KAT2A and KAT2B but no effect on H3K9ac levels, potentially due to semi-redundant functions of these HATs (24). **B**-**E**, Quantification of live Cas9(+) H3.3K27M mutant cells transduced with control, *KAT2A,* or *KAT2B* sgRNAs revealing a growth defect due to *KAT2A* and *KAT2B* KO in SU-DIPGVI cells (**B**), but no significant differences in SU-DIPGXIIIP*, BT869, or SU-DIPGXIII cell growth (**C**-**E**). **F**, Immunoblot analysis of whole cell lysates from SU-DIPGVI+Cas9 cells expressing three independent *USP22* sgRNAs confirming loss of USP22 and a corresponding increase in H2A119ub with 2 out of 3 sgRNAs (sg3 and sg4). **G**, Live SU-DIPGVI+Cas9 cell counts following transduction with control and *USP22* sgRNAs revealing no significant change in DMG growth in bulk neurosphere culture due to *USP22*KO. **H** and **I**, Average cell viability from CellTiter-Glo assays after 7 days treatment of H3.3K27M mutant cells (green curves) or non-transformed H3 wildtype control cells (black curves) with increasing doses of CPTH2, a catalytic inhibitor of KAT2A and p300/CBP (**H**) (60), or with garcinol, a natural product inhibitor of KAT2B/p300 (**I**) (54). **J,** Immunoblot analysis of SU-DIPGXIII whole cell lysates collected after 3 days of treatment with vehicle (DMSO) or CPTH2 showing a dose-dependent reduction of both H3K9ac and H3K27ac. **K** and **L**, Average number of live SU-DIPGXIII (**K**) and BT245 (**L**) cells assessed by trypan blue exclusion counted following 7 or 3 days treatment with the indicated doses of KAT2A/2B inhibitors. **M** and **N**, Dose curve analyses of H3.3K27M DMG (blue curves) and control cells (black curves) following 7 days treatment with the WDR5 WIN site inhibitor OICR9429 (**M**), or the WDR5 WBM site inhibitor WM586 (**N**), indicating a lack of selectivity for DMG cells. **O**, Heatmap of IC_50_ values from dose curve analyses using various SAGA/ATAC-targeting tool compounds across H3 wildtype and control cells, H3 wildtype HGG cells, and H3K27M mutant DMG cells highlighting selective DMG growth inhibition with GSK4027, WDR5-IN-4, and WDR-IN-6, and targeting of both H3K27M DMG and H3 wildtype HGG with USP22i-S02.

**Supplementary Figure 3.**
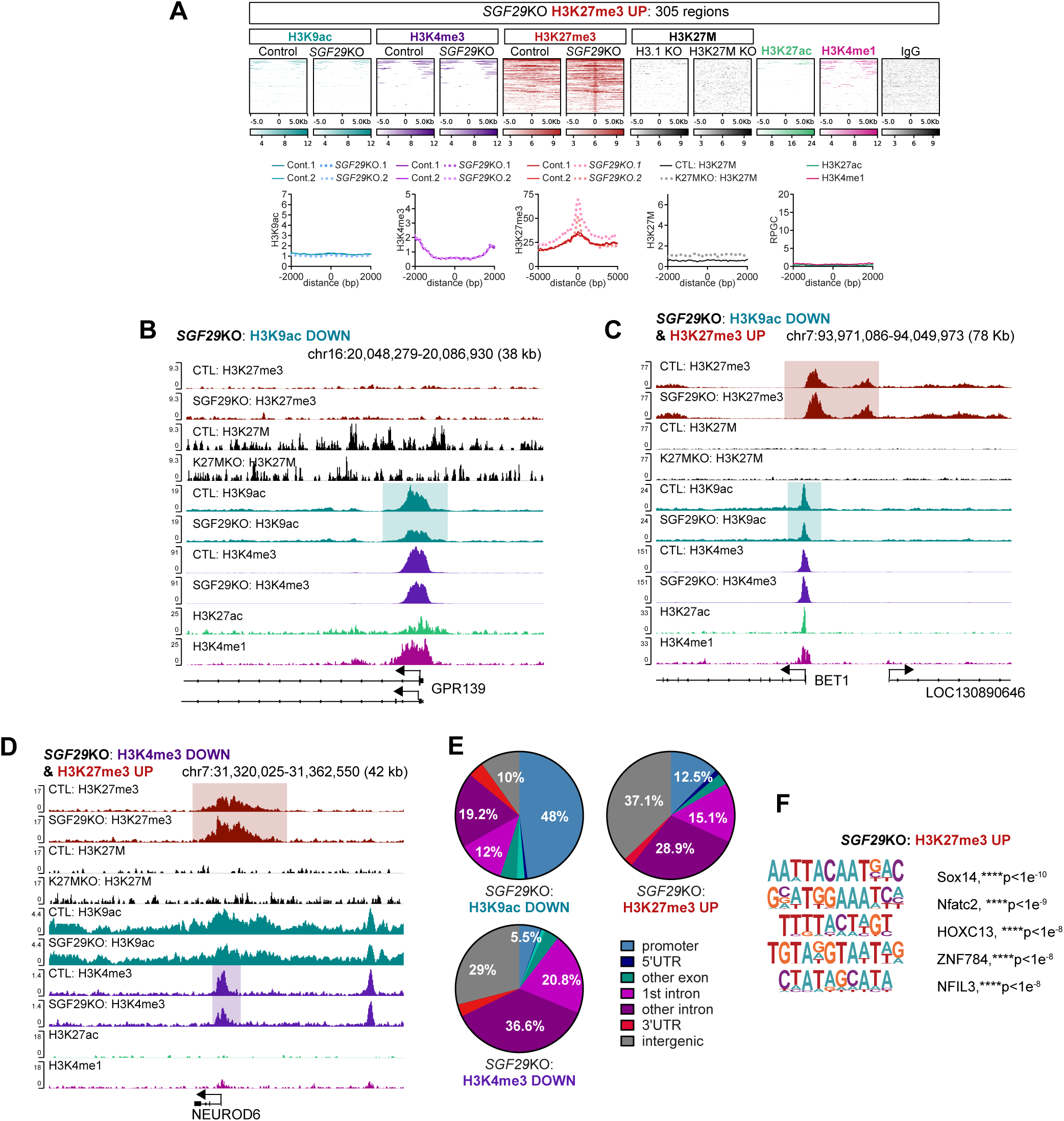
*SGF29*KO leads to repressed chromatin states at a subset of genes. **A**, Heatmap and profile plots showing H3K27me3-up peaks (305 regions) in *SGF29*KO versus control SU-DIPGXIII cells. Aligning additional ChIP-seq and chromatin profiling data at these sites reveal minimal H3K9ac, H3K4me3, H3K27M, H3K4me1, and H3K27ac binding at these regions. **B** and **C**, Genome browser snapshots showing reduced H3K9ac binding (teal boxes) at both the *GPR139* (*G Protein-Coupled Receptor 139*; **B**) and *BET1* (*Blocked Early in Transport 1 Homologue*; **C**) TSS/promoters and increased H3K27me3 at the *BET1* gene (red box; **C**) due to *SGF29*KO. **D**, Genome browser tracks at the *NEUROD6* locus showing *SGF29*KO-induced loss of H3K4me3 (purple box) and gain of H3K27me3 (red box). **E**, Pie charts showing the overlap between the H3K9ac-down, H3K4me3-down, and H3K27me3-up peaks and various genomic elements indicating that *SGF29*KO leads to reduced H3K9ac primarily at gene promoters, whereas H3K4me3 was lost and H3K27me3 was gained at both promoters and at introns/intergenic regions potentially acting as distal enhancers. **F**, Fold enrichment of transcription factor binding sites from HOMER analysis of the H3K27me3-up peaks in *SGF29*KO cells identifying binding sites for developmental transcription factors.

**Supplementary Figure 4.**
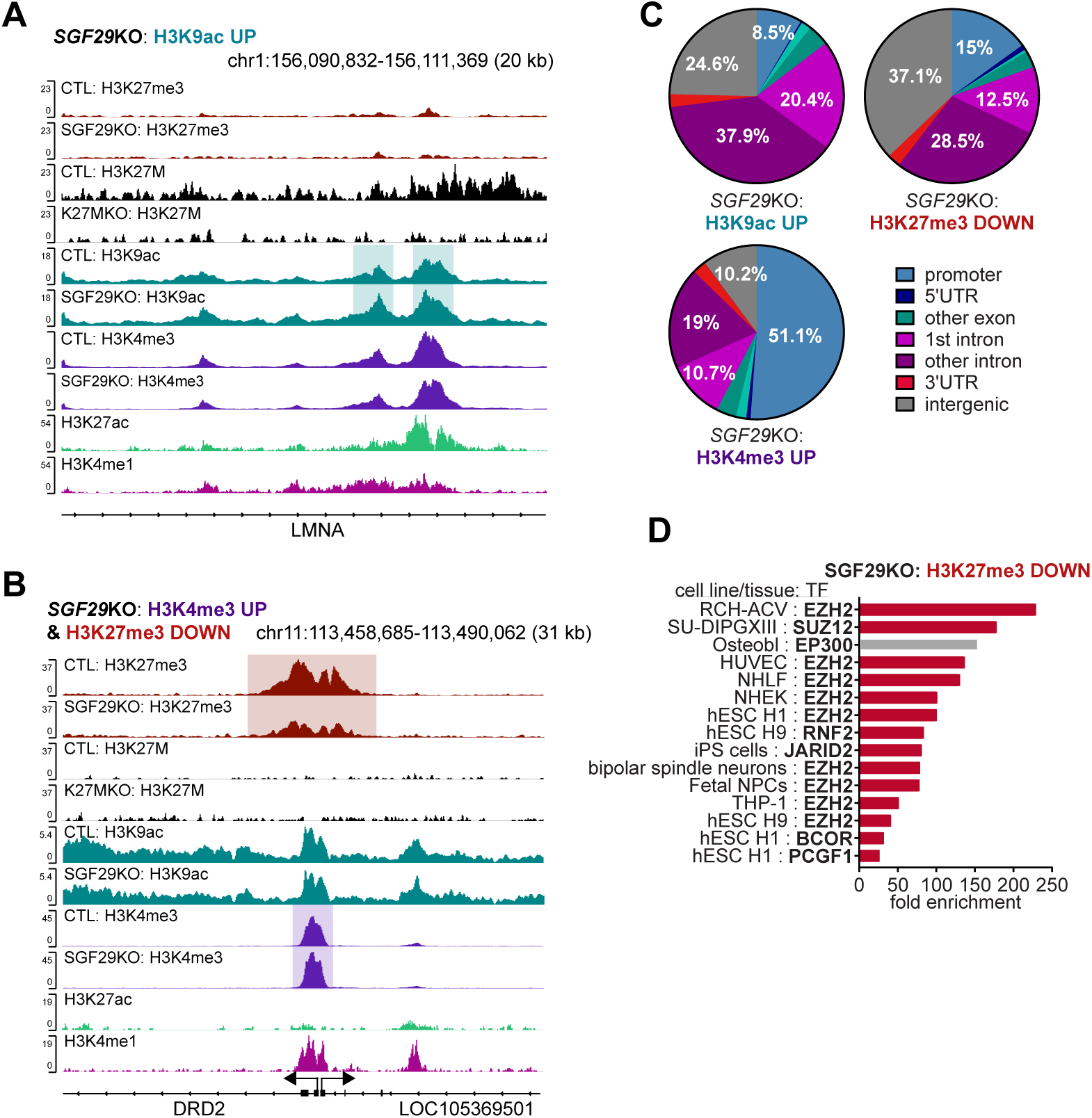
*SGF29*KO leads to active chromatin states at a subset of regions. **A** and **B**, Genome browser tracks showing increased H3K9ac (teal box) at the *LMNA* gene (**A**) and both increased H3K4me3 and decreased H3K27me3 at the DRD2 gene (**B**). **C**, Pie charts showing the overlap between the H3K9ac-up, H3K4me3-up, and H3K27me3-down peaks and various genomic elements suggesting that *SGF29*KO leads to increased H3K9ac and decreased H3K27me3 at intronic and intergenic regions and increased H3K4me3 at promoters. **D**, Fold enrichment of transcription factor binding sites, as catalogued in the ChIP ATLAS database amongst the H3K27me3-down peaks confirming an overlap with known binding sites for Polycomb-related proteins like SUZ12, EZH2, JARID2, and PCGF1.

**Supplementary Figure 5.**
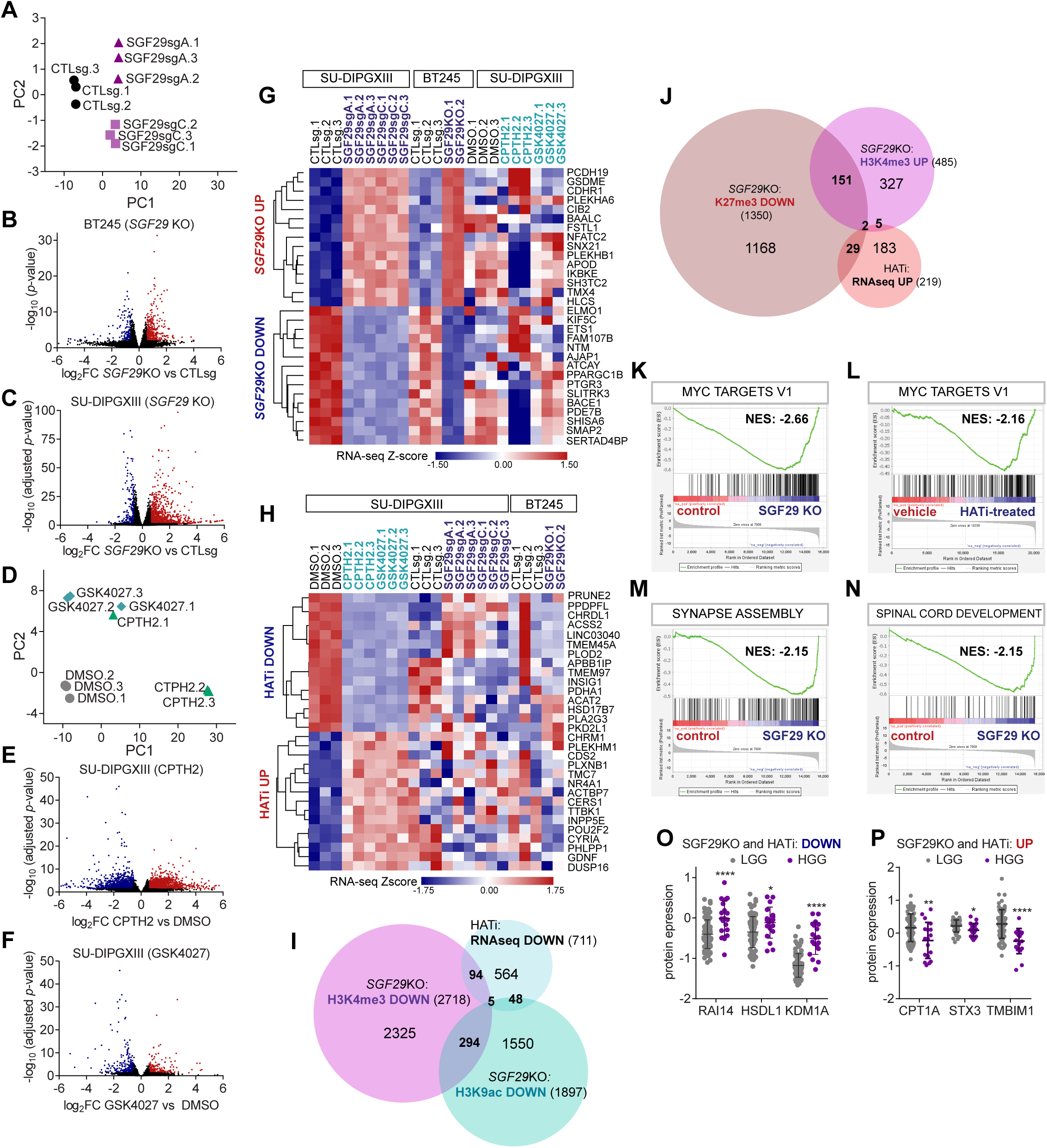
SAGA/ATAC inhibition disrupts DMG chromatin and transcriptional regulation. **A**, Principal Component Analysis (PCA) of RNA-seq data from SU-DIPGXIII showing clustering of the control and *SGF29*KO samples. **B** and **C**, Volcano plots showing log_2_ fold change and -log_10_ p-values for DEGs from *SGF29*KO versus control BT245 (**B**) and SU-DIPGXIII cells (**C**). **D**, PCA of RNA-seq data from SU-DIPGXIII showing clustering of samples treated with vehicle (DMSO),10 µM CTPH2, or 10 µM GSK4027 for 25 hours. **E** and **F**, Volcano plots showing log_2_ fold change and -log_10_ adjusted p-values for DEGs from RNA-seq analysis of CPTH2- versus vehicle-treated (**E**), or from GSK4027- versus vehicle-treated SU-DIPGXIII cells (**F**). **G** and **H**, Heatmaps of RNA-seq Z-scores summarizing DEGs observed uniquely in the *SGF29*KO condition (**G**) or following HATi treatment (**H**). **I**, Venn diagram showing overlap between HATi-downregulated transcripts and genes exhibiting H3K9ac and/or H3K4me3 loss upon *SGF29*KO. **J,** Venn diagram showing partial concordance between the genes upregulated by HATi treatment and the presence of H3K4me3-up and H3K27me3-down CUT&RUN peaks. **K**-**N**, Pre-ranked GSEA results showing that MYC target gene transcripts are reduced by both *SGF29*KO and by HATi treatment (**K** and **L**) and that transcripts associated with synaptogenesis and spinal cord development were reduced by *SGF29*KO (**M** and **N**). **N** and **O**, Protein expression from the CPTAC proteomics database of primary childhood brain tumors revealing increased (**N**) or decreased (**O**) expression of SAGA/ATAC-regulated transcripts in HGG versus LGG (see also Fig. **6A**, bold text).

**Supplementary Figure 6.**
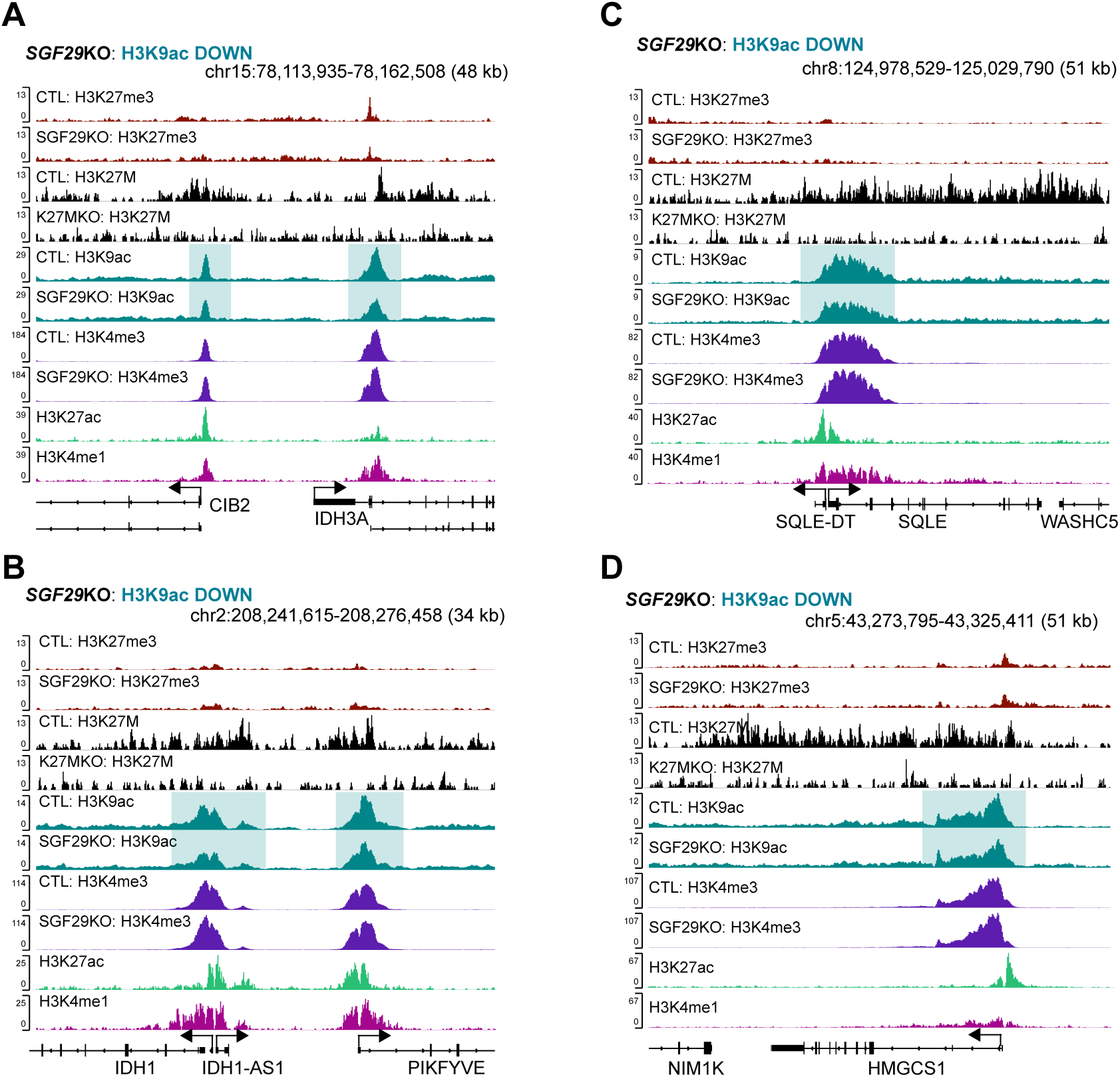
SGF29 maintains H3K9ac at the promoters/TSS of metabolic regulators. **A-D,** Genome browser snapshots of the *IDH3A* (*Isocitrate Dehydrogenase 3;* **A**), *IDH1/PIKFYVE* (*Isocitrate Dehydrogenase 1/Phosphoinositide Kinase*, *FYVE*; **B**), *SQLE* (Squalene Epoxidase; **C**), and *HMGCS1* (*HMG-CoA Synthase*; **D**) genes showing H3K27M occupancy and reduced H3K9ac (teal boxes) in *SGF29*KO versus control SU-DIPGXIII cells.

**Supplementary Figure 7.**
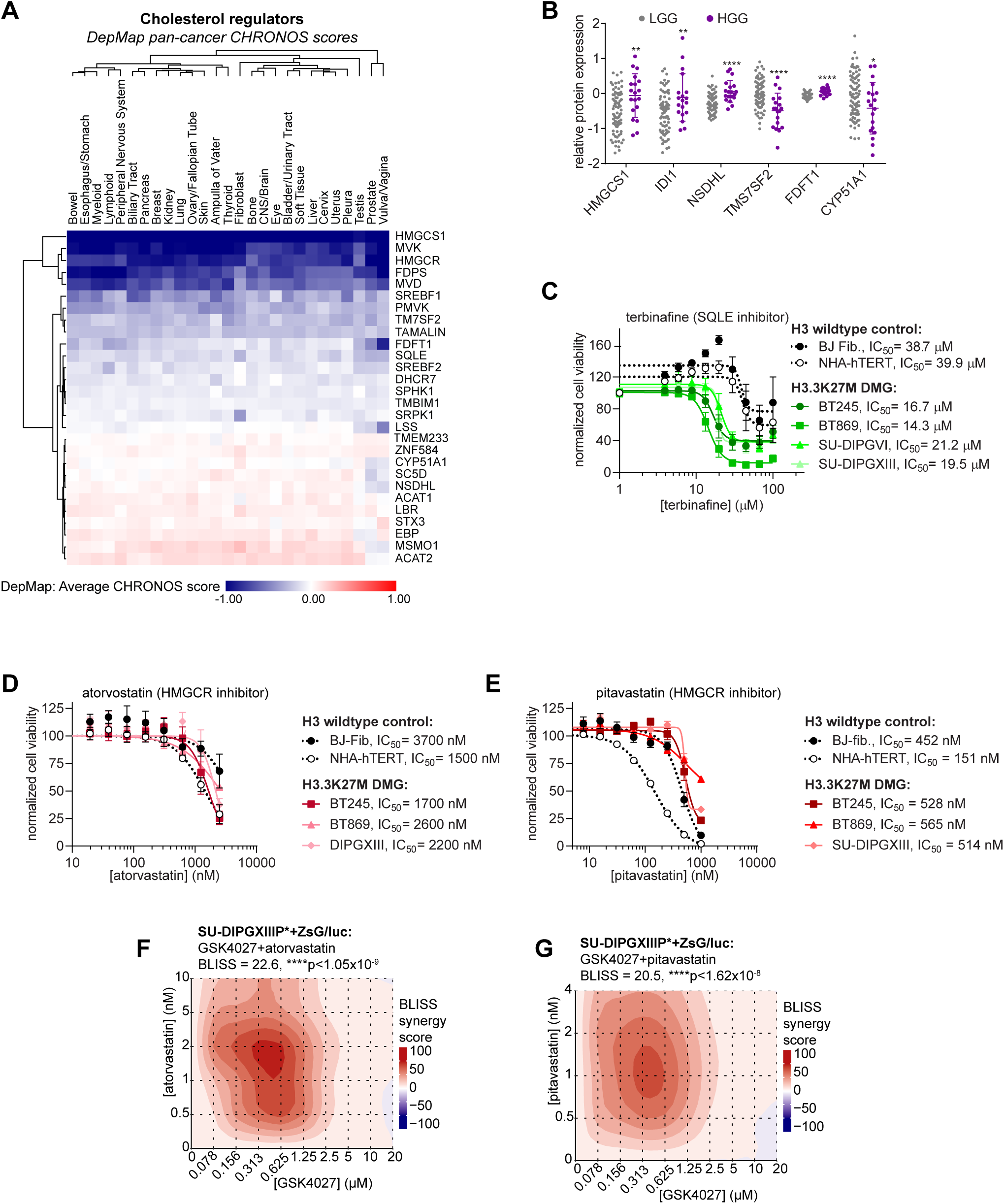
Crosstalk between SAGA/ATAC-dependent chromatin regulation and metabolic dysfunction in DMG. **A**, Heatmap summarizing average DepMap CHRONOS scores for cholesterol regulators and mevalonate pathway genes across adult cancers grouped according to tissue of origin identifying *HMGCS1*, *HMGCR*, *MVK*, *FDPS*, and *MVD* as pan-cancer dependencies. **B**, Analysis of CPTAC pediatric glioma proteomics data revealing increased expression of cholesterol synthesis regulators and mevalonate pathway genes in HGG versus LGG. **C**-**E**, Normalized CellTiter-Glo viability of H3.3K27M mutant DMG (green/red curves) versus H3 wildtype control cells (black curves) after 7 days treatment with the SQLE inhibitor terbinafine (**C**), or with the HMGCR inhibitors atorvastatin (**D**) and pitavastatin (**E**). **F** and **G**, Heatmaps of drug synergy BLISS scores from analysis of SU-DIPGXIIIP*+ZsGreen/luciferase cells responses to combinations of GSK4027 and either atorvastatin (**F**) or pitavastatin (**G**).

